# miRNome profiling of lung cancer metastases revealed a key role for miRNA-PD-L1 axis in the modulation of chemotherapy response

**DOI:** 10.1101/2022.11.23.517634

**Authors:** Roberto Cuttano, Tommaso Colangelo, Juliana Guarize, Elisa Dama, Maria Pia Cocomazzi, Francesco Mazzarelli, Valentina Melocchi, Orazio Palumbo, Elena Marino, Elena Belloni, Francesca Montani, Manuela Vecchi, Massimo Barberis, Paolo Graziano, Andrea Pasquier, Julian Sanz-Ortega, Luis M. Montuenga, Cristiano Carbonelli, Lorenzo Spaggiari, Fabrizio Bianchi

**Author notes:** Corresponding Author: Fabrizio Bianchi, Unit of Cancer Biomarkers, Institute for Stem Cell Biology, Regenerative Medicine and Innovative Therapies (ISBReMIT), Fondazione IRCCS Casa Sollievo della Sofferenza, Viale Padre Pio 7, 71013 San Giovanni Rotondo, Italy;. These authors contributed equally to this work (Tommaso Colangelo, Juliana Guarize). Non-coding RNAs and RNA-based Therapeutics, Istituto Italiano di Tecnologia, CMP^3^VdA, Via Lavoratori Vittime del Col du Mont 28, 11100 Aosta, Italy.

## Abstract

Locally-advanced non–small-cell lung cancer (NSCLC) is frequent at diagnosis and requires multimodal treatment approaches. Neoadjuvant chemotherapy (NACT) followed by surgery is the treatment of choice for operable locally-advanced NSCLC (Stage IIIA). However, the majority of patients are NACT-resistant and shows persistent lymph nodal metastases (LNmets) and an adverse outcome. Therefore, the identification of mechanisms and biomarkers of NACT resistance is paramount for ameliorating prognosis of patients with Stage IIIA NSCLC. Here, we investigated the miRNome and transcriptome of chemo naïve LNmets collected from patients with Stage IIIA NSCLC (N=64). We found that a microRNA signature accurately predicts NACT response. Mechanistically, we discovered a miR-455-5p/PD-L1 regulatory axis which drives chemotherapy resistance, hallmarks metastases with active IFN-*γ* response pathway (an inducer of PD-L1 expression), and impacts T cells viability and relative abundances in tumor-microenviroment (TME). Our data provides new biomarkers to predict NACT response and adds molecular insights relevant for improving the management of patients with locally-advanced NSCLC.

## BACKGROUND

Lung cancer is frequently diagnosed as advanced stage disease (Stage III-IV) with metastases spread to regional and distant organs in more than two-third of cases[1]. Despite the progress made in early diagnosis and treatment, the prognosis of patients remains poor with 5-year survival rates ranging from 32% to 6%, depending on the presence of regional or distant metastases, respectively[1]. One-third of patients with non-small cell lung cancer (NSCLC), i.e. the most common type of lung cancer (∼80-85% of cases), are diagnosed with locally advanced disease (Stage III). Stage III disease is heterogenous both for tumor size (from <3cm, T1; to >7cm, T4) and metastatic spreading (i.e., regional lymph nodes, N2-N3; ipsilateral peribronchial and/or ipsilateral hilar lymph nodes and intrapulmonary nodes, N1)[2]. Stage IIIA-N2 disease is prevalent and, when resectable, is preferentially treated by neoadjuvant chemotherapy (NACT; platinum-based doublet (P-doublet)) before surgery to target nodal metastases and reduce/eradicate metastatic disease. Indeed, NACT is an effective treatment in N2 patients improving the overall survival by 5% at 5-years[3]. However, clinical responses to NACT differ widely, ranging from patients achieving a complete eradication of all nodal metastases at the time of surgery (pN0) to patients having persistent metastatic disease (pN+) [4–6], which suggests the presence of different molecular features among and within nodal metastatic lesions, as recently described also in other studies [7,8]. Recently, the combination of immune checkpoint inhibitors (ICI) targeting the PD-1/PD-L1 axis (i.e., Nivolumab) with P-doublet chemotherapy in the neoadjuvant setting, showed an improved clinical management of patients with resectable NSCLC[9] and gained approval by Food and Drug Administration (FDA). In addition, other ongoing clinical trials are also evaluating the efficacy of ICI alone or in combination with NACT for stage IIIA-N2 NSCLC patients[10]. Nevertheless, the current scant knowledge of the molecular biology of metastases makes it difficult to search for cancer driver mechanisms alongside the development of predictive biomarkers and new druggable targets.

Here, by exploring the miRNA-mRNA transcriptional network of lung cancer lymph node metastases in stage IIIA-N2 disease, we derived miRNA signatures predictive of NACT response. Importantly, using *in vitro* and *in vivo* lung cancer models, we showed for the first time the role of miR-455-5p in mediating chemotherapy resistance and immune evasion by means of PD-L1 expression regulation.

## RESULTS

### Lung metastatic cells exhibit a distinct miRNA profile according to their sensitivity to NACT

We initially investigated the molecular profile of tumor metastatic cells from mediastinal lymph nodes (i.e., LNmets; station 4 and 7; see method) collected by endobronchial ultrasound transbronchial needle aspiration (EBUS-TBNA) before NACT in treatment naïve stage IIIA patients who had a complete pathological response (pN0; n=5) or with persistent disease (pN2; n=7) after P-doublet NACT (i.e., EBUS-samples; Table 1). LNmets were expanded in cell culture (Fig. 1A) as we previously showed[11]; morphological examination together with immunofluorescence staining using anti-pan-cytokeratin antibody (Pan-CK) confirmed their epithelial origin (Fig. 1B). Yet, LNmets were enriched in the expression of typical markers of cells constituting the airway epithelium (*NKX2-1*, *KRT5*, *CC10*, *SOX2*, *SFTPC*; Fig. 1C). Next, we performed high-throughput microRNA expression profiling of LNmets by TaqMan Low-density Array (TLDA; see Methods) and we detected a total of 197 miRNAs (Cqn <30.01 in at least 50% of samples for group; see Methods) (Fig. 1D-E; Data File 1). Overall, many miRNAs were downregulated in patients who developed pN2 disease (n=87, 44.9%; FC <0.67) (Fig. 1F), with 16 miRNAs (aka, LN-signature) statistically significant (p<0.05) (Fig. 1F-G). TLDA analysis of LNmets in a second independent FFPE cohort of stage III patients (n=52) collected by mediastinoscopy (i.e., MED-samples; Table 2; Fig. S1A; see Methods) resulted in the detection of 170 miRNAs (Fig. S1B), largely overlapping with those identified in EBUS-samples (Fig. S1C) and with a comparable expression level (Fig. S1D). Again, we observed a general loss of miRNA expression in patients who developed pN2 disease (Fig. S1E-F). Unsupervised clustering analysis using the LN-signature discriminated pN0 from pN2 also in this independent cohort of patients (Fig. 2A), while partial responder patients (pN1), in line with their intermediate phenotype, resulted to be scattered along the cluster (Fig. 2A). Notably, MED-samples showed a similar epithelial cell content as in EBUS-samples though with a stronger expression of markers of tumor microenvironment (TME) (*CDH5*, *PTPRC aka CD45,* and *ACTA2*) (Fig. 2B) which, on the contrary, were absent in pure epithelial LNmets (EBUS-samples). Yet, 12 out of the 16 miRNAs of the original LN-signature were also found differentially expressed in MED-samples (pN2 vs. pN0; p<0.05) (Fig. 2C and Fig. S1G). Ridge-penalized logistic regression using the LN-signature (16-miRNA model) resulted in a perfect separation of responders and non-responders in the EBUS-cohort when used as training set, which slightly decreased in the MED-cohort used as validation set (AUC=0.76) (Fig. 2D-E, Table S1). When only miRNAs detected in MED-samples were used (14-miRNA model), the model reached an AUC=0.82 in the validation set (Fig. 2D and F, Table S1). Lastly, as small numbers of biomarkers are easier to use in the clinical practice, we applied LASSO regression which identified a signature of 4 miRNAs (4-miRNA model) with an AUC of 0.81 in the validation set (Fig. 2D and G, Table S1). Importantly, the clinical model alone, built by combining all available clinical and pathological parameters, showed an AUC of 67% in the validation set which increased up to 82% when combined to miRNA-based risk models (Table 3). Collectively, these results showed a distinct pattern of miRNA expression in LNmets which is predictive of chemotherapy response.

**Fig. 1.**
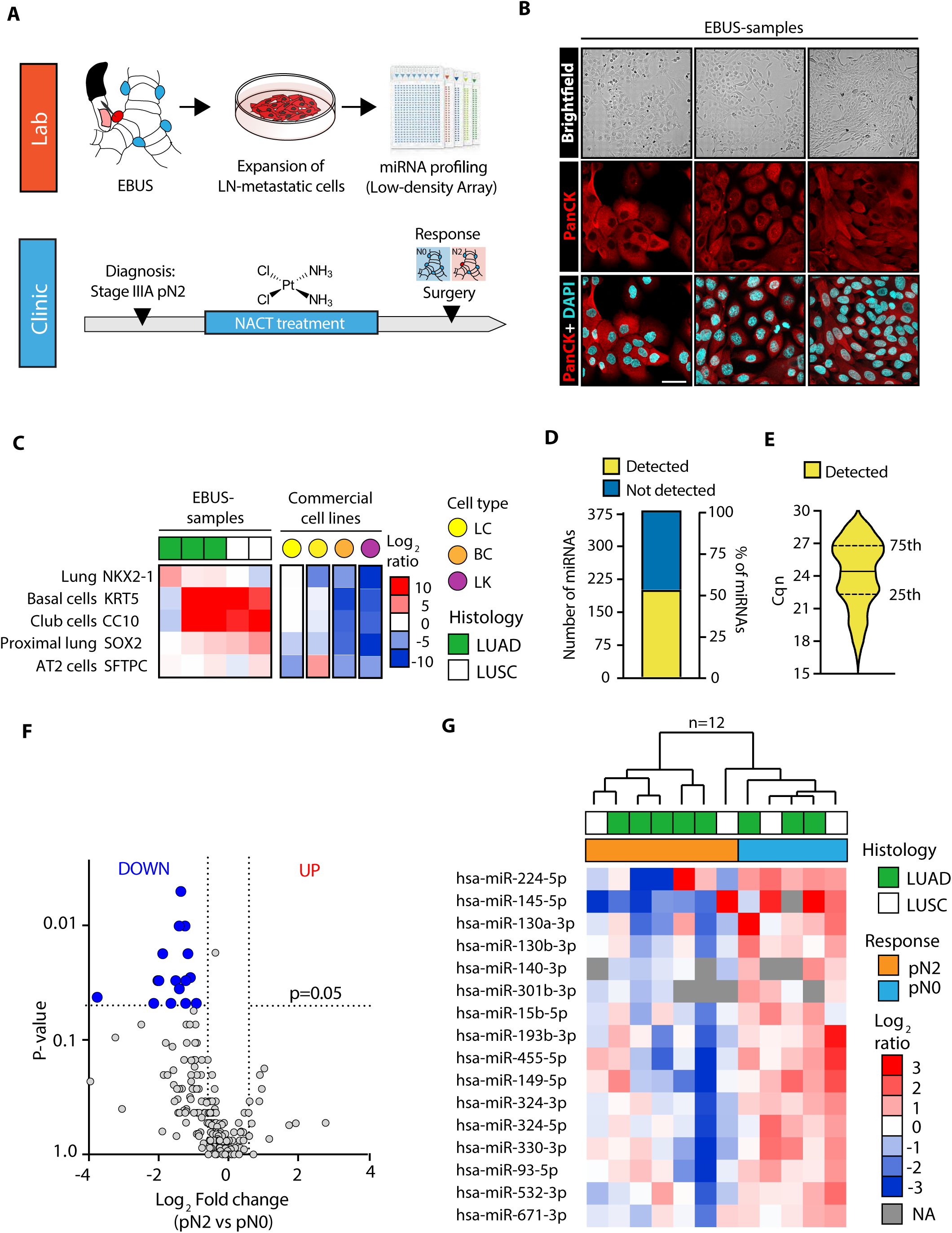
miRNA-expression profiling of LNmets collected by EBUS-TBNA. (**A**) Strategy used for miRNA-expression profiling of LNmets NSCLC cells (EBUS-samples). (**B**) *Upper panels:* brightfield images of three representative primary LNmets cell lines obtained as described in (A). Scale bar, 100 µm. *Lower panels:* representative confocal analysis of Pan-Citokeratins (PanCK) in LNmets cell lines. Pan-Citokeratins (red) identifies epithelial cells; DAPI (light blue) visualizes nuclei. Scale bar: 50µm. (C) Heat map showing qRT-PCR results of airway cell markers in five individual LN-metastatic cell lines. Two commercial lung cancer cells (LC; yellow) established from LNmets of stage IIIA NSCLC patients (NCI-H2023 and NCI-H1993) were used as positive controls for airway markers expression, while the breast cancer cells (BC; orange) MDA-MB-231 and leukemic cells (LK; magenta) HL-60 were used as negative controls. Data are log_2_-ratio. (**D**) Bar plot showing the number and percentage of miRNAs detected (yellow) or not detected (blue) in EBUS-samples. (**E**) Violin plot showing expression levels (Cqn) of all miRNAs detected in EBUS-samples. (**F**) Volcano plot showing differentially expressed miRNAs in chemoresistant (pN2; N=7) vs. chemosensitive (pN0; N=5) LNmets. Grey dot, unchanged; Blue dot, downregulated (p<0.05); Red dot, upregulated (p<0.05); Statistical significance was calculated using the Mann-Whitney U test. (**G**) Hierarchical clustering analysis of differentially expressed miRNAs (N=16, aka LN-signature) in pN2 vs pN0 LNmets. Data are log_2_-ratio. LUAD, lung adenocarcinoma; LUSC, lung squamous cell carcinoma.

**Fig. 2.**
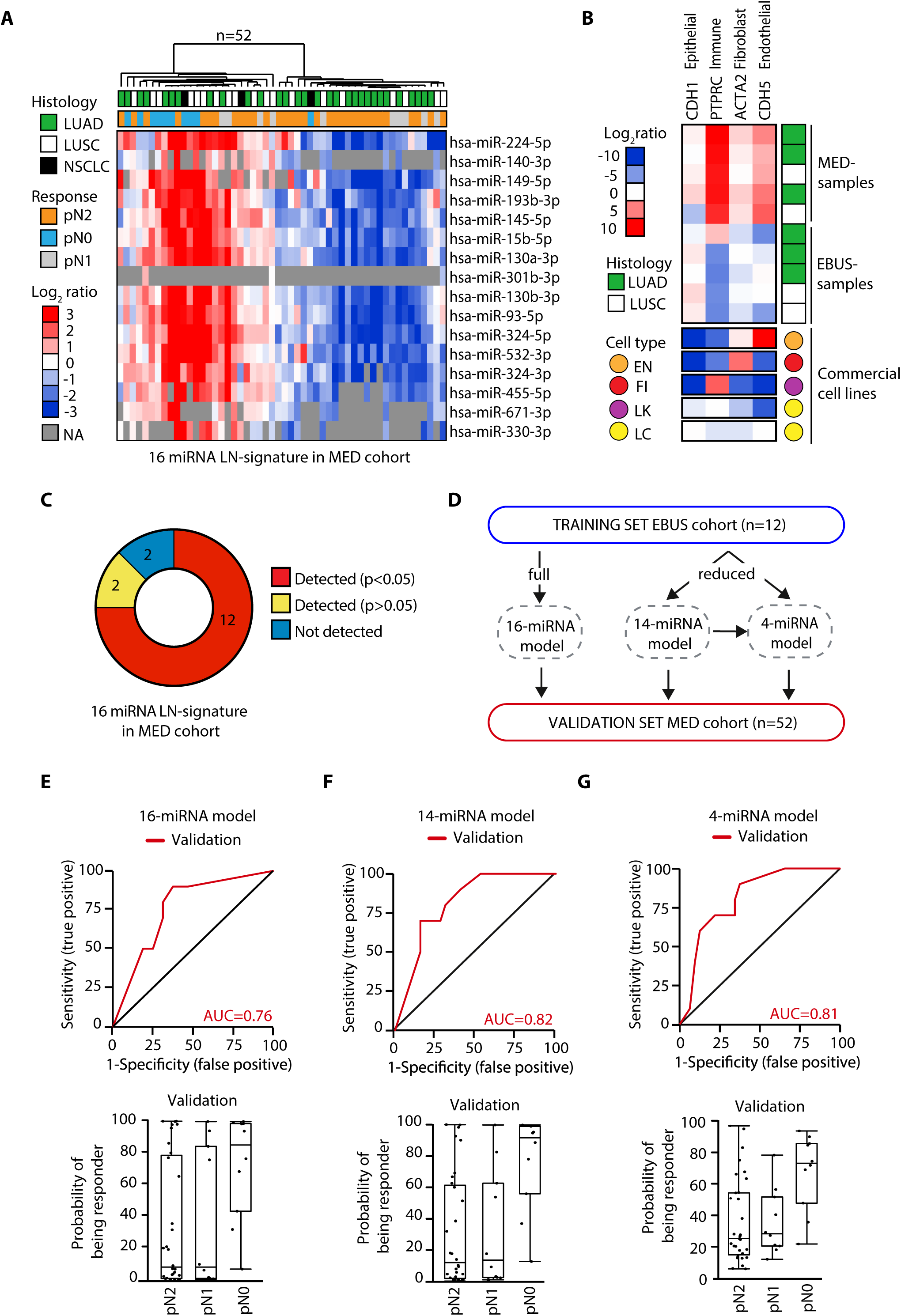
LN-signature predicts chemotherapy response of chemo-naïve lung metastatic tumor tissue collected by mediastinoscopy. (**A**) Hierarchical clustering analysis of the LN-signature in MED-samples. Data are log_2_-ratio. LUAD, lung adenocarcinoma; LUSC, lung squamous cell carcinoma; NSCLC, other non-small cell lung subtypes; NA, no available data. (**B**) Heat map showing gene expression of the indicated marker analyzed by qRT-PCR in LNmets (EBUS-samples, N=5; and MED-samples, N=5). NCI-H2023 and NCI-H1993 lung cancer cells (LC, yellow) were used as positive controls for the expression of epithelial marker while HUVEC (EN, orange), WI38 (FI, red) and HL-60 cells (LK, magenta) were used as positive control for endothelial, fibroblast and immune-like markers expression, respectively. Data are log_2_-ratio. (**C**) Pie chart showing the number of miRNAs of LN-signature (N=16) that were found differentially expressed between pN0 and pN2 samples in MED-cohort. (**D**) Schematic representation of strategy adopted to derive miRNA-based NACT predictive models. (**E** to **G**) *Upper panels:* receiver operating characteristic (ROC) curves of the 16-miRNA model (E), 14-miRNA model (F) and 4-miRNA model (G) in the validation set (MED-samples, red). *Lower panels*: box plot of the predicted probability of being a responder according to the 16-miRNA model (E), 14-miRNA model (F) and 4-miRNA model (G).

**Table 1.** Clinical-pathological characteristics of EBUS cohort. Abbreviations: LUAD, Lung adenocarcinoma; LUSC, Lung squamous cell carcinoma; NACT, Neoadjuvant chemotherapy; CDDP, Cisplatin; GEM, Gemcitabine; CBDCA, Carboplatin; NA, no available data.

|  | **ALL**<br>**(n=12)** | **pN0 (n=5)** | **pN2 (n=7)** |
| --- | --- | --- | --- |
| **Age [years]** |  |  |  |
| Median (Q1;Q3) | 67 (62;72) | 69 (66;74) | 64 (57;72) |
| **Gender** |  |  |  |
| Female | 5 (42%) | 2 (40%) | 3 (43%) |
| Male | 7 (58%) | 3 (60%) | 4 (57%) |
| **Histology** |  |  |  |
| LUAD | 8 (67%) | 3 (60%) | 5 (71%) |
| LUSC | 4 (33%) | 2 (40%) | 2 (29%) |
| **Stage** |  |  |  |
| IIIA | 12 (100%) | 5 (100%) | 7 (100%) |
| **NACT regimen** |  |  |  |
| CDDP+GEM | 10 (83%) | 5 (100%) | 5 (71%) |
| CBDCA+GEM | 1 (8%) | 0 (0%) | 1 (14%) |
| NA | 1 (8%) | 0 (0%) | 1 (14%) |
| **Number of NACT cycles** |  |  |  |
| 3 cycles | 8 (67%) | 2 (40%) | 6 (86%) |
| 4 cycles | 3 (25%) | 3 (60%) | 0 (0%) |
| NA | 1 (8%) | 0 (0%) | 1 (14%) |

*Percentages could not add up to 100% due to rounding*

**Table 2.** Clinical-pathological characteristics of MED cohort. Abbreviations: LUAD, Lung adenocarcinoma; LUSC, Lung squamous cell carcinoma; NSCLC, other non-small cell lung subtypes; NACT, Neoadjuvant chemotherapy; CDDP, Cisplatin; GEM, Gemcitabine; CBDCA, Carboplatin; VNR, Vilnorebine; NA, no available data.

|  | **ALL (n=52)** | **pN0 (n=10)** | **pN2 (n=32)** | **pN1 (n=10)** |
| --- | --- | --- | --- | --- |
| **Age** |  |  |  |  |
| Median (Q1;Q3) | 62.12 (59.18; 68.24) | 62.05 (59.81; 70.32) | 61.32 (58.82; 66.97) | 62.95 (59.86; 66.32) |
| **Gender** |  |  |  |  |
| Female | 15 (28.8%) | 3 (30%) | 9 (28.1%) | 3 (30%) |
| Male | 37 (71.2%) | 7 (70%) | 23 (71.9%) | 7 (70%) |
| **Histology** |  |  |  |  |
| LUAD | 30 (57.7%) | 4 (40%) | 22 (68.7%) | 4 (40%) |
| LUSC | 19 (36.5%) | 4 (40%) | 9 (28.1%) | 6 (60%) |
| NSCLC | 3 (5.8%) | 2 (20%) | 1 (3.1%) | 0 (0%) |
| **Stage** |  |  |  |  |
| IIIA | 46 (88.5%) | 8 (80%) | 29 (90.6%) | 9 (90%) |
| IIIB | 6 (11.5%) | 2 (20%) | 3 (9.4%) | 1 (10%) |
| **NACT regimen** |  |  |  |  |
| CDDP+ALIMTA | 6 (11.5%) | 0 (0%) | 5 (15.6%) | 1 (10%) |
| CDDP+GEM | 41 (78.8%) | 9 (90%) | 25 (78.1%) | 7 (70%) |
| CDDP+TAXOTERE | 1 (1.9%) | 0 (0%) | 1 (3.1%) | 0 (0%) |
| CDDP+VNR | 4 (7.7%) | 1 (10%) | 0 (0%) | 2 (20%) |
| VNR+GEM | 1 (1.9) | 0 (0%) | 1 (3.1%) | 0 (0%) |
| **Number of NACT cycles** |  |  |  |  |
| 2-3 cycles | 42 (80.8%) | 9 (90%) | 24 (75%) | 9 (90%) |
| 4-5 cycles | 9 (17.3%) | 1 (10%) | 7 (25%) | 1 (10%) |

*Percentages could not add up to 100% due to rounding*

**Table 3.** Combination of clinical model with miRNA predictive model in MED cohort. (**A**) Performance of single predictive models based on clinical information (age, gender or histology) or miRNA expression (16, 14 and 4 miRNAs). (**B**-**D)** Combination of clinical models with 16 miRNA risk score (B), 14 miRNA risk score (C) and 4 miRNA risk score (D). Odds Ratio (OR), P-value calculated by Wald Test and AUC of indicated models are reported in the table.

| MED-cohort (validation set) | OR (95% CI) | Wald p-value | AUC |
| --- | --- | --- | --- |
| **clinical model** |  |  |  |
| Age (5-unit increase) | 1.04 (0.65-1.66) | 0.88 | 67% |
| Gender (male vs. female) | 0.58 (0.10-3.31) | 0.54 |  |
| Histology (LUSC/NSCLC NOS vs. LUAD) | 3.88 (0.79-19.18) | 0.1 |  |
| **miRNA model** |  |  |  |
| 16-miRNA risk score | 1.20 (1.00-1.44) | 0.046 | 76% |
| 14-miRNA risk score | 1.37 (1.07-1.75) | 0.0125 | 82% |
| 4-miRNA risk score | 2.00 (1.17-3.42) | 0.0116 | 81% |

(B)
| MED-cohort (validation set) | OR (95% CI) | Wald p-value | AUC |
| --- | --- | --- | --- |
| **clinical model** |  |  |  |
| Age (5-unit increase) | 1.12 (0.67-1.86) | 0.67 | 77% |
| Gender (male vs. female) | 0.46 (0.07-2.98) | 0.42 |  |
| Histology (LUSC/NSCLC NOS vs. LUAD) | 2.71 (0.51-14.48) | 0.24 |  |
| **miRNA model** |  |  |  |
| 16-miRNA risk score | 1.19 (0.99-1.45) | 0.07 |  |

(C)
| MED-cohort (validation set) | OR (95% CI) | Wald p-value | AUC |
| --- | --- | --- | --- |
| **clinical model** |  |  |  |
| Age (5-unit increase) | 1.13 (0.67-1.93) | 0.88 | 82% |
| Gender (male vs. female) | 0.40 (0.06-2.78) | 0.54 |  |
| Histology (LUSC/NSCLC NOS vs. LUAD) | 2.40 (0.42-13.79) | 0.1 |  |
| **miRNA model** |  |  |  |
| 14-miRNA risk score | 1.37 (1.06-1.77) | 0.0175 |  |

(D)
| MED-cohort (validation set) | OR (95% CI) | Wald p-value | AUC |
| --- | --- | --- | --- |
| **clinical model** |  |  |  |
| Age (5-unit increase) | 1.14 (0.67-1.94) | 0.63 | 80% |
| Gender (male vs. female) | 0.46 (0.07-3.11) | 0.42 |  |
| Histology (LUSC/NSCLC NOS vs. LUAD) | 1.95 (0.33-11.38) | 0.46 |  |
| **miRNA model** |  |  |  |
| 4-miRNA risk score | 1.98 (1.11-3.54) | 0.0216 |  |

### Functional analysis of predictive microRNAs to NACT response

We then used the LN-signature to identify mechanisms of chemotherapy resistance. First, we analyzed public drug screening datasets, such as CTRPv2, GDSC1-2 and PRISM [12–16], to retrieve cisplatin (i.e., the backbone component of NACT) sensitivity data in NSCLC cell lines for which miRNA expression data were available (CCLE dataset). Unexpectedly, cytotoxic effect of cisplatin was negligible in the majority of the cell lines at the indicated doses (Fig. 3A, Table S2). However, we noticed that, at least in the GDSC2 dataset, DMSO was used as compound vehicle, which is known to rapidly inactivate cisplatin[17]. Therefore, we performed a small-scale drug screening to test cisplatin sensitivity (dissolved in NaCl 0.9%) of a panel of metastatic NSCLC cell lines. Cells were treated with increasing doses of cisplatin and drug sensitivity was measured by sigmoidal curve fitting (Fig. 3B). NSCLC cell lines exhibited a heterogenous sensitivity profile to cisplatin, with potency (IC_50_) ranging from 1.5 to 11µM and efficacy (E_max_) calculated at the peak plasma concentration of cisplatin upon injection (Cmax, ∼12µM: [18,19]) from 0 to 0.5 relative cell viability (Fig. 3C). When we analyzed the expression of our LN-signature in chemo-naïve NSCLC cell lines, we observed a variable degree of association between IC_50_/E_max_ values and miRNAs expression (Fig. 3D). Interestingly, miR-455-5p was the top scoring in terms of negative correlation to cisplatin IC_50_/Emax values (IC_50,_ r=-0.82 p=0.034; E_max_, r=-0.71 p=0.088) (Fig. 3D-E). As shown above, this was in line with the downregulation of miR-455-5p observed in LNmets of NACT-resistant patients (Fig. 3F). We also scored a negative correlation for miR-140-3p (IC_50,_ r=-0.76 p=0.037; E_max_, r=-0.69 p=0.069) whose overexpression was indeed shown to sensitize NSCLC cells to cisplatin [20,21] (Fig. 3D).

**Fig. 3.**
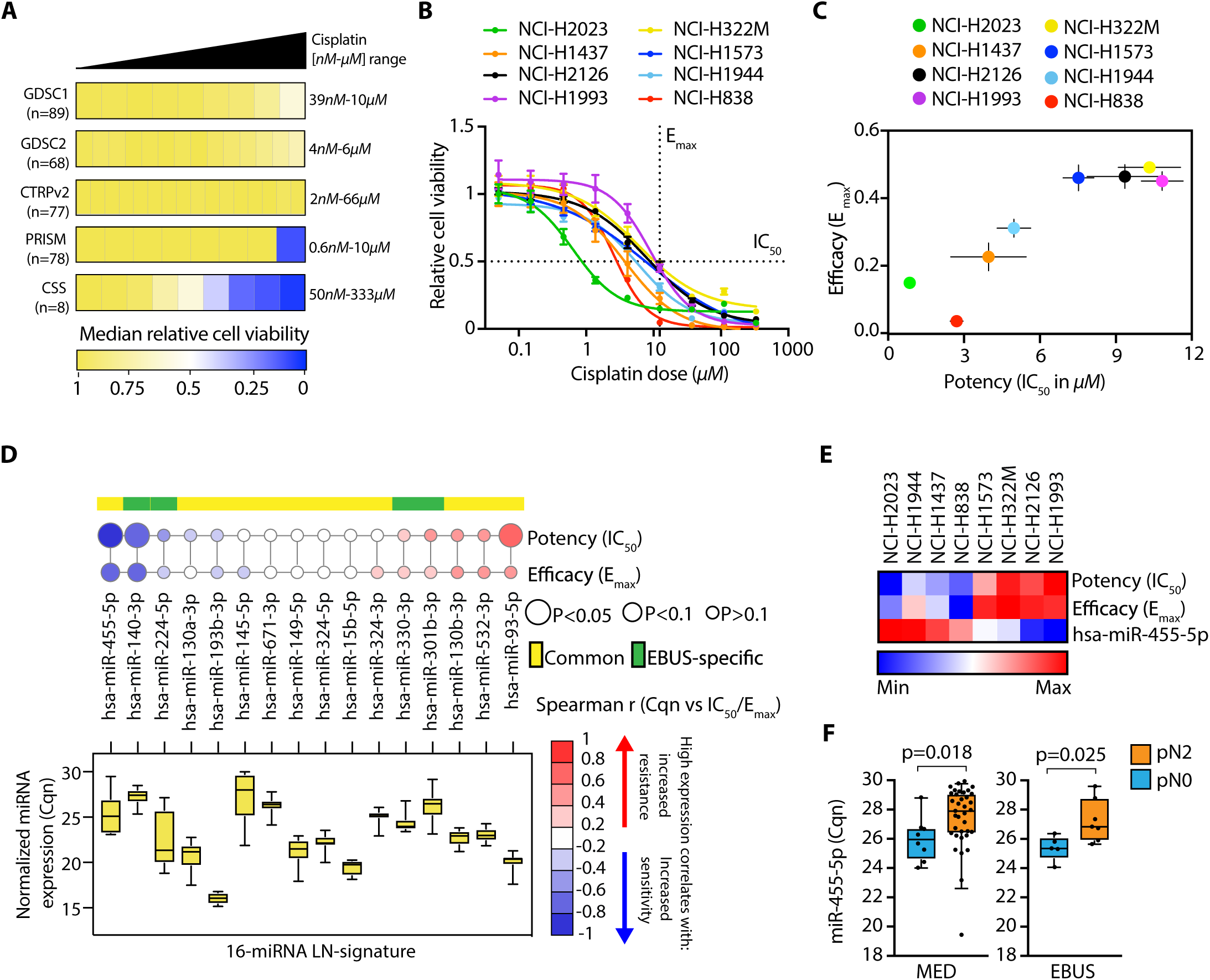
Basal levels of miR-455-5p negatively correlate with cisplatin resistance *in vitro*. (**A**) Heatmap of cell viability values (median normalized) of NSCLC cell lines at increasing concentrations of cisplatin. Heatmap square represents an individual drug concentration drug. For each dataset, number of cell lines and concentration range used (minimum-maximum) are indicated. (**B**) Dose-response curves of the indicated NSCLC cell lines treated with cisplatin for 72 hours. Error bars indicate SEM (N=3 to 5). (**C**) Distribution of potency (IC_50_) versus efficacy values (E_max_) of cisplatin in the indicated NSCLC cell lines. Data are mean ± SEM (N=3 to 5). (**D**) *Upper panel*: bubble plot reporting correlation coefficient (r) between basal level of normalized miRNA expression (Cqn) and IC_50_ or E_max_ values. The size of the bubble is proportional to statistical significance calculated by the Spearman correlation test, while colors indicate *r* coefficient. Yellow: common differentially expressed miRNAs in both EBUS- and MED-samples; Green: miRNAs differentially expressed in EBUS-samples only. *Lower panel*: box plot representing the expression levels (Cqn) of miRNAs in the panel of NSCLC cell lines. (**E**) Heatmap of mean value of IC_50_, E_max_ and miR-455-5p expression (Cqn) in the indicated cell lines. (**F**) Box plot showing miR-455-5p expression levels (Cqn) in chemoresistant (pN2) and chemoresponsive (pN0) patients in MED- and EBUS-samples. P-values were calculated by the Mann-Whitney U test.

### miR-455-5p regulates cisplatin resistance of lung metastatic cells

Next, we investigated whether miR-455-5p was sufficient to modulate chemotherapy response of NSCLC cells. To this end, we took advantage of the NCI-H1993 cell line which *i)* was derived from LNmets of a stage IIIA NSCLC patient, *ii)* is a miR-455-5p low expressing cell line and *iii)* has a higher resistance to cisplatin (Fig. 3E). NCI-H1993 cells were transfected with a miR-455-5p mimic (OE) or a negative mimic control (CTRL) and the increased levels of miR-455-5p after overexpression were confirmed by qRT-PCR (Fig. 4A). Importantly, we observed that miR-455-5p OE in NCI-H1993 strongly increased sensitivity to cisplatin (Fig. 4B) with a significant decrease of cisplatin potency in comparison to CTRL cells (Fig. 4C). We then investigated whether miR-455-5p could play a role also in acquiring cisplatin resistance and thus we treated the cisplatin sensitive NCI-H2023 cell line (Fig. 3C) with increasing doses of cisplatin during cycles of drug on (4 days) and drug off (1–2 weeks) (Fig. 4D). Long-term treatment with cisplatin resulted in the generation of a resistant variant of the NCI-H2023 cell line namely the NCI-H2023-CDDP-R (aka, CDDP-R), which was characterized by a significant increase in both IC_50_ and E_max_ in comparison to parental cells (Fig. S2A-B). The acquirement of resistance to cisplatin was accompanied by the acquisition of a typical elongated cell shape (Fig. S2C), an increased mRNA and protein expression of epithelial-to-mesenchymal transition (EMT) master regulators (i.e., ZEB1, SLUG and TWIST1) and of mesenchymal/stem cells markers (VIM, ACTA-2, CD90) (Fig. S2D-F)[22]. Indeed, the gene expression profiling of parental and CDDP-R cells (Fig. S2G) followed by gene set enrichment analysis (GSEA) using ‘Hallmark genes set’ collection, revealed that the “EMT gene signature” was the highest one significantly enriched in cisplatin resistant cells (Fig. S2H-I, Table S3A). Lastly, we observed a reduced proliferation rate and a higher migratory/invasive capability of CDDP-R cells (Fig. S2J-L).

**Fig. 4.**
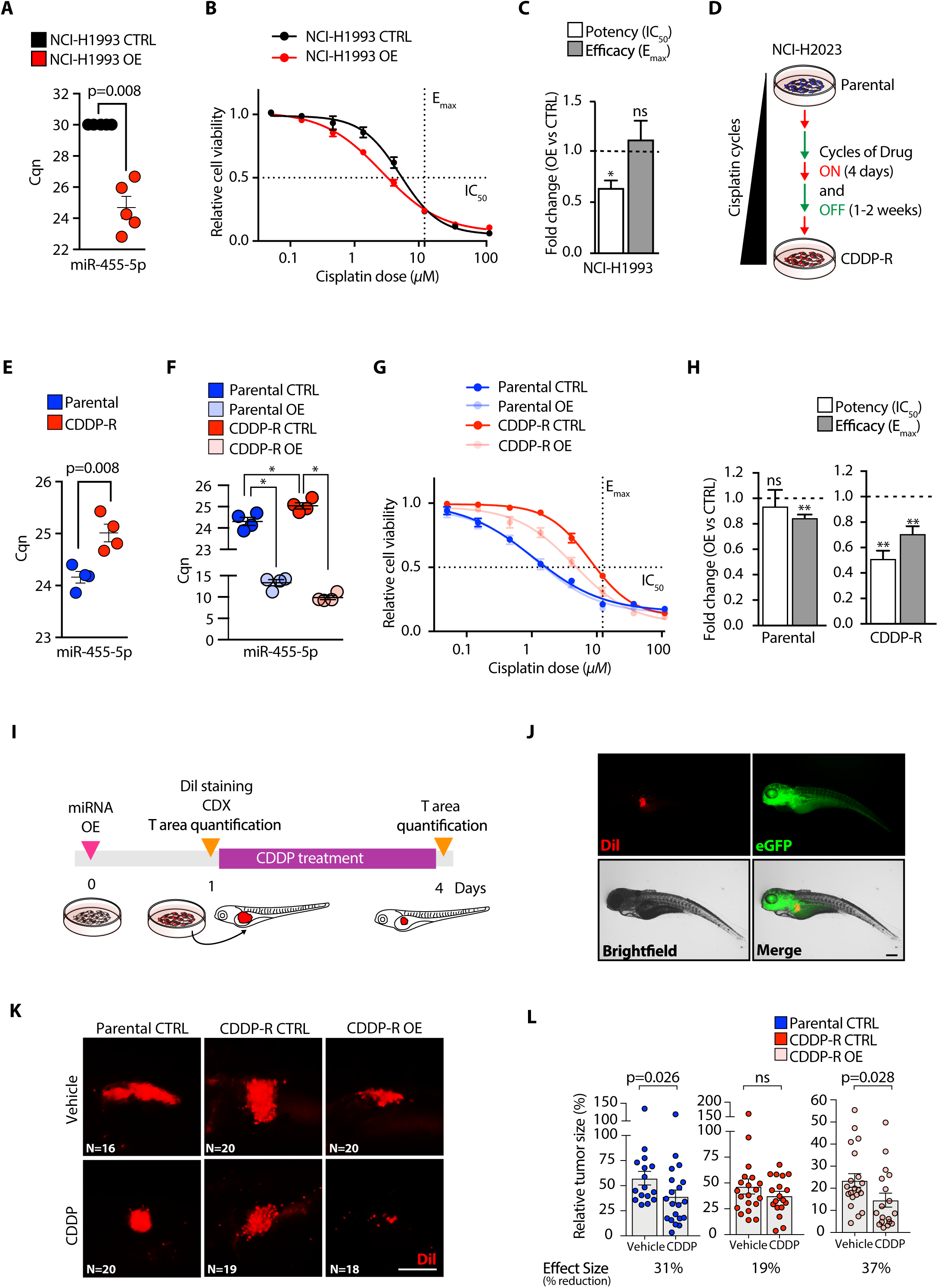
miR-455-5p modulates cisplatin resistance *in vitro* and *in vivo*. (**A**) qRT-PCR of miR-455-5p in NCI-H1993 transfected with a miR-455-5p mimic (NCI-H1993 OE) or a negative control mimic (NCI-H1993 CTRL). Data, expressed as normalized Cq (Cqn), are mean ± SEM (N=5). P-value was calculated by the Mann-Whitney U test. (**B**) Dose-response curves of NCI-H1993 CTRL and NCI-H1993 OE cells treated with cisplatin for 72 hours. Error bars indicate SEM (N=4). (**C**) Bar plot of cisplatin potency and efficacy of NCI-H1993 CTRL and NCI-H1993 OE cells. Data are mean ± SEM (N=4). Fold change is relative to NCI-H1993 CTRL. P-value was calculated by one sample t-test. *P<0.05; ns, not significant. (D) Generation of a model of *in vitro* acquired cisplatin resistance. (**E**) qRT-PCR of miR-455-5p in Parental and CDDP-R cell lines. Data, expressed as Cqn, are mean ± SEM (N=4). P-value was calculated by t-test with Welch’s correction. (**F**) qRT-PCR of miR-455-5p in Parental and CDDP-R transfected either with a miR-455-5p mimic (i.e., Parental OE, and CDDP-R OE) or a negative control mimic (i.e., Parental CTRL, and CDDP-R CTRL). Data, expressed as Cqn, are mean ± SEM (N=4). P-value was calculated using the Mann-Whitney U test. *P <0.05. (**G**) Dose-response curves of indicated cell lines treated with cisplatin for 72 hours. Error bars indicate SEM (N=5). (**H**) Bar plot of cisplatin potency and efficacy of Parental CTRL, Parental OE, CDDP-R CTRL and CDDP-R OE cells. Data are mean ± SEM (N=5). Fold change is relative to CTRL. P-value was calculated by one sample t-test. **P <0.01; ns, not significant. (**I**) Schematic representation of zCDX model to monitor chemotherapy response *in vivo*. (**J**) Representative fluorescence images of zebrafish larvae injected with tumor cells. Dil (red) identifies tumor cells; eGFP (green) visualizes blood vessels. Scale bar: 200µm. (**K**) Representative fluorescence images of tumor masses upon 3 days of cisplatin treatment. Dil (red) identifies tumor cells. Scale bar: 200µm. (**L**) Size distribution of tumor masses derived from indicated cell lines. Columns represent mean ± SEM (N=16-20, for each condition). Results are shown as relative tumor size (i.e. percent change in tumor size by comparing day 4 vs. day 1). Effect size is expressed as percent reduction in mean value of tumor size. P-value were calculated by the Mann-Whitney U test.

In line with the above observations, miR-455-5p was significantly downregulated after acquisition of cisplatin resistance in CDDP-R vs. parental cells (Fig. 4E). We, therefore, transfected miR-455-5p in both parental and CDDP-R cells (Fig. 4F) and performed cell viability analysis upon cisplatin treatment (Fig. 4G). Strikingly, miR-455-5p overexpression in CDDP-R cells induced cisplatin sensitivity both in terms of potency and efficacy when compared to parental cells or parental cells overexpressing miR-455-5p (Fig. 4G-H), thus suggesting a specific miR-455-5p-addiction in resistant cells.

We validated such findings also *in vivo* by using a zebrafish cell derived xenograft (zCDX) model which was recently shown to be valuable in oncology research [23,24]. First, parental and CDDP-R cells overexpressing miR-455-5p or not, as a control, were fluorescently labeled and then injected into the perivitelline space of zebrafish larvae (Fig. 4I). qRT-PCR analysis confirmed miR-455-5p OE before cell inoculation (Fig. S3A). Next, zebrafish embryos were treated with cisplatin at a dose near to Cmax (∼16µM) and tumor growth analyzed (Fig. 4I-J). Implantation rate was 100% in both cell lines upon injection (at day 0), with parental cells that formed slightly smaller tumors when compared to tumors formed by CDDP-R cells (Fig. S3B-C). The cisplatin treatment induced a significant reduction in the tumor size of the parental tumors but not of the CDDP-R ones (Fig. 4K-L). Strikingly, miR-455-5p overexpression re-sensitized CDDP-R tumors to cisplatin (Fig. 4K-L). Yet, miR-455-5p OE alone caused a significant reduction of the tumor burden in CDDP-R untreated resistant tumors (Fig. 4K-L). This is in line with *in vitro* data where miR-455-5p OE impaired tumor cell proliferation (Fig. S4A-B) and with the observation that high miR-455-5p expressing tumors from TGCA-LUAD cohort are smaller in size when compared to low miR-455-5p ones (Table S4).

### PD-L1 is a direct molecular link between miR-455-5p and cisplatin resistance

We then asked which molecular mechanisms can be influenced by miR-455-5p and their role in cisplatin resistance. To tackle this, we reconstructed miRNA-mRNA transcriptional networks by performing transcriptome analysis of LNmets (MED-samples) which identified 1702 differentially expressed genes (DEGs) (fold change>|1.5|; p<0.05) in pN2 vs. pN0 patients (Fig. 5A). GSEA using a curated gene set representing miR-455-5p predicted target genes (n=349, Data File 3; see Methods) revealed a positive enrichment (FDR<0.05) of miR-455-5p targets in pN2 patients which was coherent with previously observed loss of miR-455-5p expression (Fig. 5B). Next, we used the ‘Hallmark genes set’ collections in GSEA which revealed a number of pathways involved in the regulation of proliferation, metabolism, immune evasion, development and response to cellular stresses, enriched in LNmets of pN2 patients (FDR<0.05) (Fig. 5C, Table S3B). To functionally dissect regulation of pN2-enriched pathways, we transfected NCI-H1993 and CDDP-R cells with a miR-455-5p mimic (OE) or a negative mimic control (CTRL) and performed transcriptome analysis. GSEA confirmed the modulation of miR-455-5p target genes upon miRNA overexpression (Fig. 5B). Strikingly, comparative analysis of significantly enriched ‘Hallmark gene sets’ (FDR<0.05) in MED-samples and in the two NSCLC cell lines (NCI-H1993 and CDDP-R) revealed that ‘INTERFERON-ALPHA (IFN-α) RESPONSE’ and ‘INTERFERON-GAMMA (IFN-*γ*) RESPONSE’ were overlapping and enriched in LNmets of pN2 patients likewise in low-miR-455-5p expressing NSCLC cell lines with the same trend of regulation (Fig. 5C-E, Table S3B-D). Next, we looked among genes belonging to IFN-α and IFN-*γ* response pathways to search for putative miR-455-5p target genes by TargetScan analysis[25]. *BATF2*, *CMPK2*, *IRF2*, *MYD88*, *SOCS3* and *PD-L1* (aka *CD274*) genes were all predicted to be targeted by miR-455-5p (Fig. 5F). Among these genes, PD-L1 expression was previously reported to be found increased after NACT treatment in NSCLC. [26–28]. Moreover, besides the well-known role of PD-L1 in the regulation of T cell activity through the interaction with the receptor PD-1, it was also found to regulate critical functions of cancer cells in a cell autonomous way, including chemotherapy resistance [29,30]. Therefore, we speculated that miR-455-5p regulation would impact chemotherapy response through PD-L1 direct regulation. Overall, we analyzed PD-L1 expression (mRNA, total and cell-surface protein) in our panel of NSCLC cell lines (Fig. S5A-C) and found that a higher expression of PD-L1 was associated with cisplatin resistance (Fig. S5D-E). Furthermore, when we silenced PD-L1 expression by siRNAs in NCI-H1993 cells the sensitivity to cisplatin increased significantly (Fig. S5F-H). Conversely, the acquisition of cisplatin resistance was accompanied by a concomitant increase of PD-L1 expression in CDDP-R when compared to parental cells (Fig. S6A-C). Accordingly, silencing of PD-L1 by siRNAs in CDDP-R cells (Fig. S6D) was able to strongly enhance cisplatin sensitivity when compared to control cells (Fig. S6E-F), whilst no effect was scored in the parental cell lines where PD-L1 expression was low (Fig. S6E-F).

**Fig. 5.**
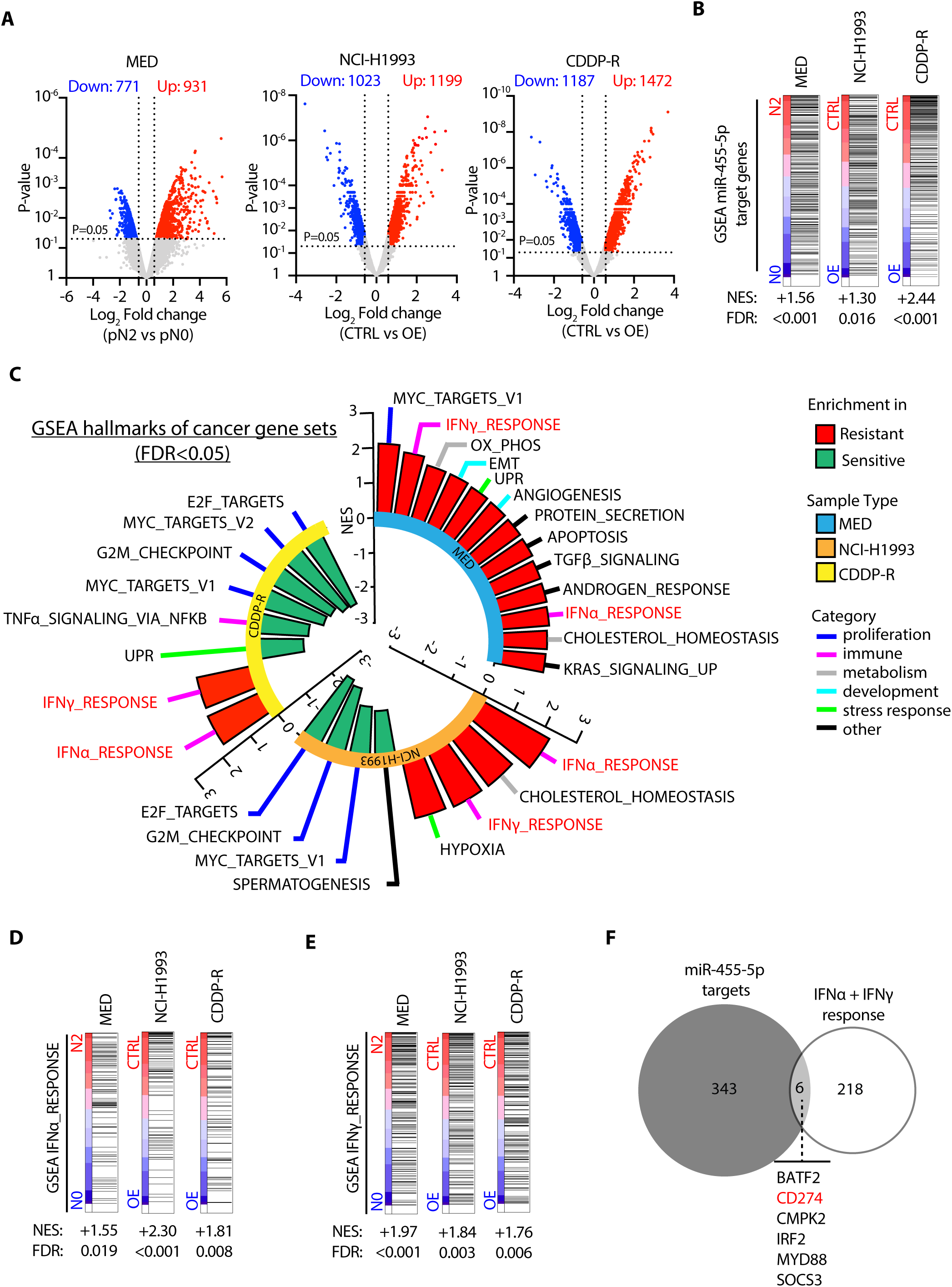
miR-455-5p modulates the expression of genes involved in interferon response. (**A**) Volcano plot showing differentially expressed genes found by microarray analysis. *Left panel:* pN2 vs. pN0 (MED-samples). *Central panel:* NCI-H1993 CTRL vs NCI-H1993 OE cells (N=2). *Right panel:* CDDP-R CTRL (N=2) vs CDDP-R OE cells (N=2). Grey dot, unchanged genes; Blue dot, downregulated genes (p-value <0.05; FC <-1.5); Red dot, upregulated genes (p-value <0.05; FC >1.5). P-value was calculated using the Limma moderated t-test. (**B**) GSEA using miR-455-5p predicted target genes in pN2 vs. pN0 (MED-samples), H1993 CTRL vs H1993 OE or CDDP-R CTRL vs CDDP-R OE. NES, normalized enrichment score; FDR, false-discovery rate. (**C**) Circular plot showing GSEA results using the ‘Hallmark gene sets’ collection in pN2 vs pN0 (MED-samples), H1993 CTRL vs H1993 OE and CDDP-R CTRL vs CDDP-R OE. In red, common enriched gene signatures having the same trend of regulation in all experimental conditions. (**D** and **E**) GSEA of (D) IFN-α and (E) IFN-γ response gene sets in pN2 vs pN0 (MED-samples), H1993 CTRL vs H1993 OE and CDDP-R CTRL vs CDDP-R OE. (**F**) Venn diagram representing the overlap of genes between IFN-α/IFN-γ response gene sets and miR-455-5p target genes.

### miR-455-5p/PD-L1 axis contributes to cisplatin resistance in lung metastatic cells

We then searched for predicted miRNA-binding sites in the 3’ untranslated region (3’-UTR) of *PD-L1* (aka *CD274*) which revealed a binding site (8-mer) for miR-455-5p (Fig. 5F and 6A). Indeed, we found an inverse correlation between miR-455-5p expression and PD-L1 protein amount in our panel of NSCLC cell lines (Fig. 6B). Yet, PD-L1 mRNA levels were found to be strongly upregulated in LNmets of pN2 (i.e., low miR-455-5p) vs. pN0 (i.e., high miR455-5p) patients (Fig. 6C). Remarkably, miR-455-5p expression and PD-L1 tumor proportion score showed a trend of inverse correlation also in primary NSCLC from two other independent cohorts of patients (the CSS and CIMA-CUN cohorts; Table S5; Fig. 6D-E). We also analyzed miRNA- and RNA-seq data from the TGCA-LUAD and TGCA-LUSC cohorts (N_LUAD_=507, N_LUSC_=473; Fig. 6F). When tumor samples were stratified based on the miR-455-5p expression level (‘High’, ‘Int’ and ‘Low’; see Methods) we observed an inverse correlation between miR-455-5p and PD-L1 expression (Fig. 6F). Lastly, we investigated miR-455-5p and PD-L1 association in a publicly available dataset of NSCLC patients after chemotherapy treatment (N=131,[31]; Fig. S8 A-C; see also Supplementary Methods). GSEA using a curated gene set representing miR-455-5p predicted target genes (n=349, Data File 3; see Methods) revealed a positive enrichment (FDR<0.05) of miR-455-5p targets in high PD-L1 chemoresistant NSCLC (Fig S8A). Notably, miR-455 gene is located within the intron of COL27A1 gene[32], thus we used COL27A1 expression as a surrogate of miR-455-5p expression as we previously showed[33]. Strikingly, we found that there was a significant negative correlation between COL72A1 and CD274 expression (Fig. S8B-C) which further corroborated that a high PD-L1 expression was usually associated to a lower miR-455-5p expression in chemoresistant NSCLC. Next, we transfected NCI-H1993 and CDDP-R cells with miR-455-5p mimic and analyzed PD-L1 expression *in vitro*: miR-455-5p OE decreased the level of cell-surface PD-L1 protein of NCI-H1993 and CDDP-R cells (Fig. 6G), while such effect was negligible in low-PD-L1 expressing parental cells (Fig. 6G). Importantly, similar results were obtained when we forced the expression of miR-455-5p in a primary LNmets cell line (i.e., the EBUS-52 cell line) established in our lab (Fig. 6G) (see Methods). To test the direct effect of miR-455-5p on PD-L1 expression regulation, we took advantage of custom-designed oligonucleotides (target site blockers; TSBs) that specifically prevent the binding of miR-455-5p to the *PD-L1* 3′-UTR. Transfection of TSBs in CDDP-R cells rescued PD-L1 loss of expression upon miR-455-5p OE (Fig. 6H). Strikingly, the rescue of PD-L1 expression upon TSB transfection resulted in the recovery of cisplatin resistance of CDDP-R/miR-455-5p OE cells (Fig. 6I), thus suggesting that miR-455-5p regulates cisplatin response in a PD-L1 dependent manner.

**Fig. 6.**
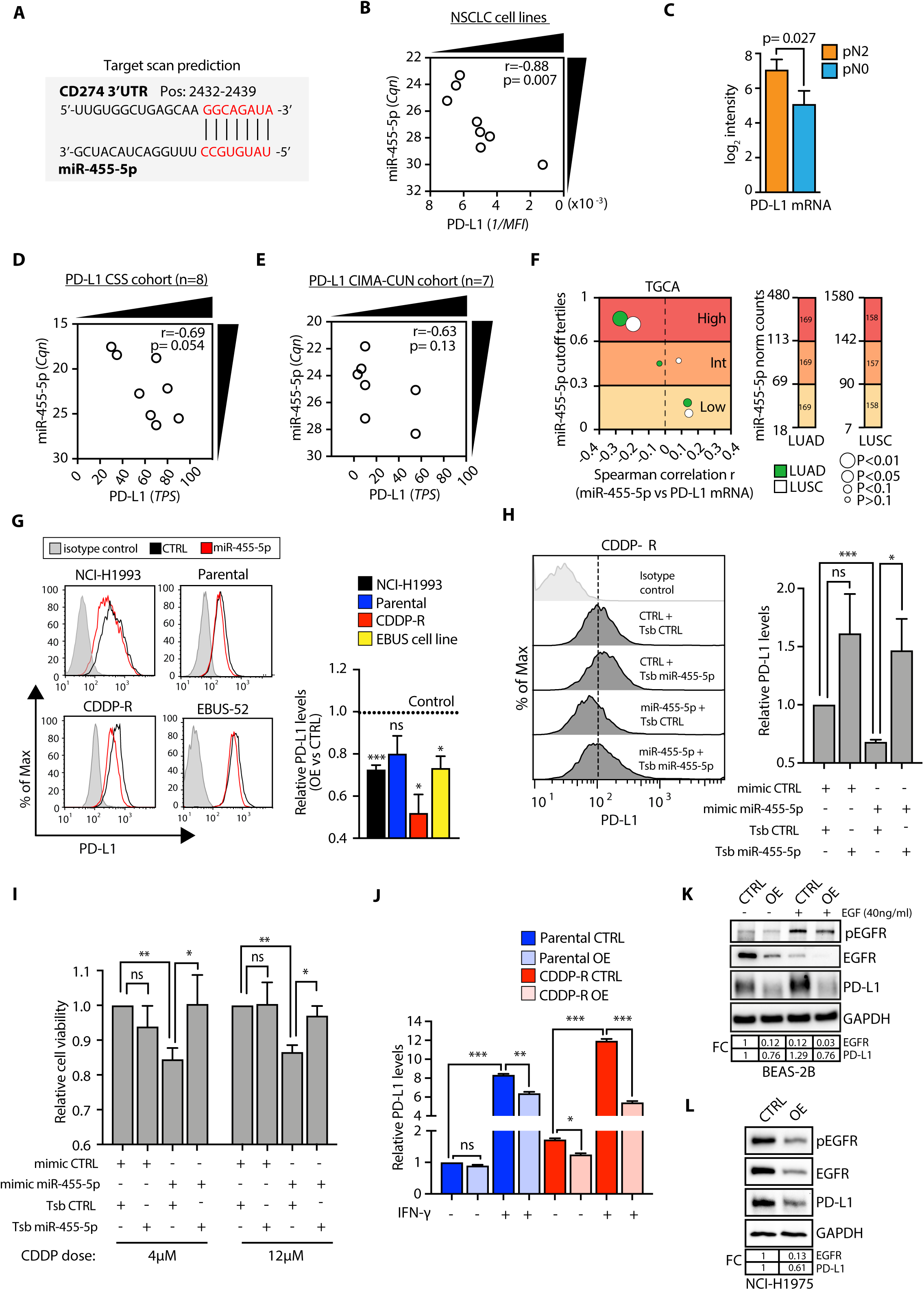
miR-455-5p regulates cisplatin resistance through direct regulation of PD-L1 expression. (**A**) Target Scan prediction of miR-455-5p binding (seed sequence in red) to human *PD-L1* 3’UTR. (**B**) Spearman correlation analysis of cell surface PD-L1 expression (reciprocal of mean fluorescence intensity values) and miR-455-5p levels (Cqn) in the panel of NSCLC cell lines. (**C**) Bar plot of *PD-L1* expression (microarray log_2_ intensity) in pN2 and pN0 patients (MED-samples). Error bars represent SEM. P-value was calculated by Limma moderated t-test. (**D** and **E**) Distribution of PD-L1 expression (TPS [tumor proportion score],) and miR-455-5p levels (Cqn) in NSCLC primary tumors obtained from CSS cohort (D) and CIMA-CUN Cohort (E). (**F**) Correlation analysis of miR-455-5p levels with PD-L1 mRNA in tumors from TGCA-LUAD and TGCA-LUSC cohorts Left: Bubble plots report the correlation coefficients. Size of the bubbles indicates statistical significance. Right: Bar plot reporting the value of miR-455-5p normalized count for each tertile threshold in TGCA-LUAD and TGCA-LUSC cohorts. The number of patients was reported inside the bar. (**G**) Representative flow cytometry histogram plots (left) and quantification (right) of PD-L1 median fluorescence intensity (MFI) in the indicated cell lines treated with a miR-455-5p mimic (OE) or a negative control mimic (CTRL). Results are shown as fold change of MFI relative to CTRL cells. Data are mean ± SEM (N=4 or 5). P-values were calculated by one sample t-test. *P<0.05, **P<0.01, ***P<0.001; ns, not significant. (**H**) Representative flow cytometry histogram plots (left) and quantification (right) of cell surface PD-L1 MFI in CDDP-R cells transfected with a miR-455-5p mimic or a negative control in the presence of a scramble TSB or a PD-L1-specific miR-455-5p TSB. Data are reported as fold change in MFI relative to CDDP-R cells transfected with a CTRL miRNA mimic and with a scramble TSB. Data are mean ± SEM (N=6). P-values were calculated by one sample t-test. *P<0.05, **P<0.001; ns, not significant. (**I**) Bar plot representing cell viability (Fold change relative to CTRL mimic in the presence of a scramble TSB) of CDDP-R cells transfected as in (G) and treated for 72h with cisplatin at the indicated doses. Data are mean ± SEM (N=5). P-values were calculated by one sample t-test. *P<0.05, **P<0.01; ns, not significant. (**J**) Bar plot representing cell-surface PD-L1 expression in the indicated cell lines stimulated for 48 hours with ± 40 ng/ml of IFN-γ. The result is shown as fold change in the MFI relative to Parental CTRL cells. Data are mean ± SEM (N=3). P-values were calculated by one sample t-test. *P<0.05, **P<0.01, ***P<0.001; ns, not significant. (**K**) Immunoblot analysis of pEGFR, EGFR and PD-L1 in BEAS-2B transfected with a miR-455-5p mimic or a negative control and treated for 36 hours with ± 40ng/ml of EGF. GAPDH was used as loading control. (**L**) Immunoblot analysis of pEGFR, EGFR and PD-L1 expression in NCI-H1975 transfected with a miR-455-5p mimic or a negative control. GAPDH was used as loading control.

In cancer, PD-L1 expression is induced upon exposure to interferons produced by activated Natural Killer (NK) and T cells in the TME [34,35]. We herein showed the enrichment of IFN-α and IFN-γ response pathways in low-expressing miR-455-5p cells and LNmets from pN2 patients (Fig. 5D-E). Thus, we asked whether miR-455-5p OE could affect IFN-mediated induction of PD-L1 expression. In line with our hypothesis, miR-455-5p OE was able to attenuate IFN-γ mediated PD-L1 upregulation both in parental and CDDP-R cells (Fig. 6J). Since PD-L1 expression in tumor cells can be influenced by the aberrant activation of oncogenic signals, such as MYC, ALK, MEK-ERK, RAS and EGFR[36], and that miR-455-5p was reported to directly regulate the EGFR expression[37], we then investigated whether miR-455-5p could interfere with the EGF mediated PD-L1 expression. Interestingly, miR-455-5p OE was able to reduce the EGFR and PD-L1 expression independently of the EGF stimulation, both in normal bronchial epithelial cells (i.e., BEASB-2B) (Fig. 6K) and in NCI-H1975 lung cancer cells (which express high levels of PD-L1 due to presence of the L858R/T790M double activating mutations of *EGFR*[38]) (Fig. 6L). Notably, miR-455-5p was also predicted to target IRF2 (Fig. 5F), a well-known transcriptional repressor of PD-L1 expression [39,40]. Indeed, we found that miR-455-5p overexpression strongly reduced IRF2 expression (Fig. S9A-B) which suggests an additional miR-455-5p/IRF2 axis potentially functioning as a regulator of miR-455-5p/PD-L1 mechanism (Fig. S9C), a possibility which warrants further investigation.

### miR-455-5p overexpression decreases T-cell apoptosis

The interaction of PD-L1 with its cognate receptor PD-1 inhibits the proliferation and activation of T cells[36]. Therefore, we asked ourselves whether miR-455-5p-dependent PD-L1 regulation in tumor cells may impact T cells viability. To this purpose, we took advantage of Jurkat cells, a leukemic T cell line widely used in the literature for T cell signaling studies[41]. NCI-H1975 cells (miR-455-5p OE or CTRL) were co-cultured for 72 hours with Jurkat cells in the presence of CD3/CD28/CD2 soluble antibody complexes to induce activation and PD-1 expression on the T cell surface (Fig. 7A). Strikingly, miR-455-5p OE decreased the percentage of apoptotic T cells when compared to T cells co-cultured with NCI-H1975 CTRL cells (Fig. 7B-C; Fig. S7A). Likewise, we observed a significant reduction of apoptotic T cells when we directly silenced PD-L1 in NCI-H1975 (Fig. 7B-C; Fig. S7A). Next, we analyzed the correlation of miR-455-5p expression with CD8 T cell infiltration in two independent cohorts of primary NSCLC tumors (the CSS and CIMA-CUN cohorts; Table S6; Fig. 7D-E). The analysis revealed a positive correlation between miR-455-5p expression and the percentage of CD8 T cells in high tumor-infiltrating lymphocytes (TILs) tumors (Fig. 7D-E). Strikingly, when we performed a pooled analysis (n=47) by combining the two cohorts, we confirmed that higher level of miR-455-5p was associated to a higher infiltration of CD8 T cells (Fig. 7 F). Furthermore, we leveraged the TCGA-LUAD and -LUSC datasets to grasp further information about CD8 T cells subsets infiltration in NSCLC samples high-/low-miR-455-5p expressing: i) TCGA samples were stratified in ‘High’, ‘Int’ and ‘Low’ miR-455-5p expressing samples (see Methods); ii) PD-L1 expression likewise expression signatures related to CD8 exhausted T cells[42] and of IFN response were analyzed in High/Int/Low miR-455-5p tumor subsets (Fig. 7G; see Methods). Strikingly, the expression levels of miR-455-5p were inversely correlated to signatures of enriched exhausted CD8+ T cell (aka GET) and of IFN response (Fig. 7G) in LUAD tumors, thus further reinforcing the link among miR-455-5p, PD-L1 and impact on T cells viability. Lastly, the analysis of the distribution of ‘Immune Subtypes’ introduced by Thorsson et al.[43] revealed, in LUAD low-miR-455-5p expressing samples, a depletion of the ‘inflammatory subtype (C3) (enriched in pro-inflammatory T helper Th1 and Th17 cells) which enhances CD8+ T cells cytotoxicity (Fig. 7G). Contrariwise, the miR-455-5p expression had no effects on the immune subtypes of LUSC tumors which, by and large, showed a distinct immune composition in comparison to LUAD tumors due to the predominance of the C2 subtype and the absence of the C3 subtype (Fig. 7G). Notably, when we analyzed N2 metastasis by the CIBERSORTx algorithm[44], we found that pN2 MED-samples were characterized by a trend in the reduction of cytotoxic cells, such as NK activated cells and T cell CD8, which was in line with our previous observations (Fig. S10A, Table S7). Moreover, pN2 and pN0 metastases were also characterized by varying expression levels of MHC and immune-inhibitors molecules (Fig. S10B).

**Fig. 7.**
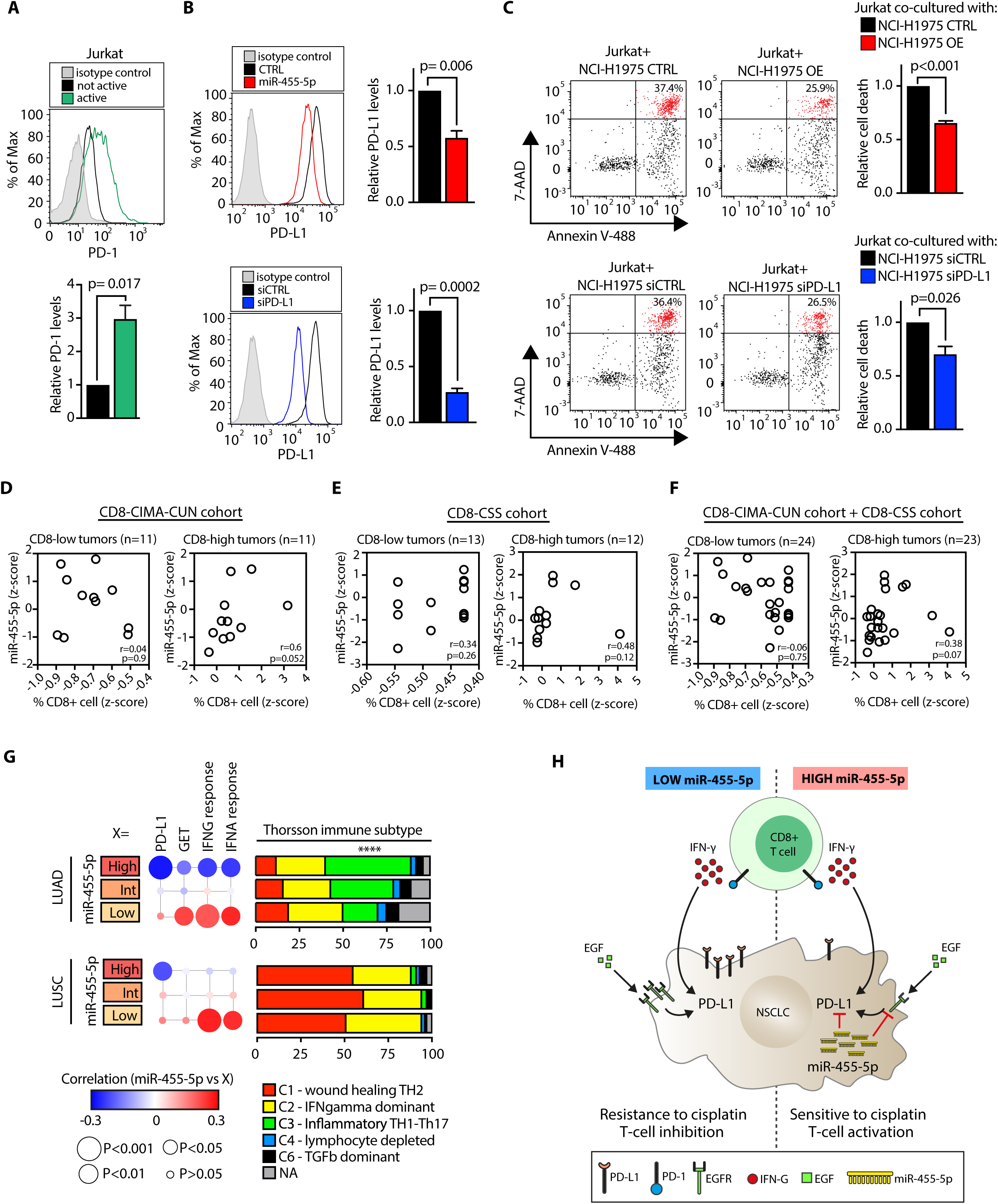
miR-455-5p overexpression decreases T cell apoptosis. (**A**) Representative flow cytometry histogram plot (upper panel) and quantification (lower panel) of PD-1 MFI in Jurkat cells stimulated either with ± CD3/CD28/CD2 soluble antibody complexes for 72 hours. Results are shown as fold change of MFI relative to not active cells. Data are mean ± SEM (N=4). P-value was calculated by one sample t-test. (**B** and **C**) NCI-H1975 cells transfected with the indicated oligos were exposed to IFN-γ for 8 hours and then co-cultured for 72 hours with Jurkat cells in the presence of T cell activator. (B) Representative flow cytometry histogram plots (left) and quantification (right) of PD-L1 MFI at the indicated experimental conditions. Results are shown as fold change of MFI relative to control conditions. Data are mean ± SEM (N=4). P-values were calculated by one sample t-test. (C) Analysis of Jurkat apoptosis rate co-cultured with the indicated cell lines by AnnexinV/7-AAD staining. Right panels: Representative flow cytometric plots (left) and quantification (right) of apoptotic dead Jurkat cells (Annexin V+, 7-AAD+; highlighted in red). Results are shown as fold change of apoptotic dead cells relative to matched control conditions. Data are mean ± SEM (N=4). P-values were calculated by one sample t-test. (**D-E-F**) Distribution of the percentage of CD8+ cells and miR-455-5p expression, expressed as z-score, in NSCLC primary tumors from CD8-CIMA-CUN (D), CD8-CSS Cohort (E) and after pooling together the two cohorts (F). Tumors were stratified in high and low CD8-tumors based on the median value of CD8+ z-score. (**G**) *Left:* correlation analysis of miR-455-5p levels with PD-L1 mRNA, gene signature for exhausted CD8+ T cell (GET), IFN-γ and IFN-α response in tumors from TGCA-LUAD and TGCA-LUSC cohorts. Bubble plots reported the correlation coefficients for miR-455-5p expression with the indicated variables. The size of the bubbles indicates statistical significance calculated by the Spearman correlation analysis. Right: Bar plot reporting the Thorsson immune subtype of TGCA-LUAD and - LUSC tumors according to miR-455-5p expression. P-value was calculated by using the t test for equality of proportions (High vs Low). ****P<0.001 (referred to C3). (**H**) Schematic model of the effects of miR-455-5p-dependent PD-L1 regulation in NSCLC.

Overall, these data suggest that miR-455-5p-dependent inhibition of PD-L1 expression may affect CD8 T cell phenotype thus improving T cell antitumor immune response.

## DISCUSSION

Patients with locally-advanced lung cancer treated by NACT in combination with surgery had a better survival than patients treated by surgery alone, in randomized trials[45]. However, response rate to NACT is still suboptimal due to the clinical and biological heterogeneity of lung tumors. Recent improvements have been made by introducing the use of ICI (e.g., nivolumab, pembrolizumab, atezolizumab; [46–48]) in combination with cisplatin-based chemotherapy, to trigger the immune response against primary and metastatic lung cancer lesions[49]. Yet, the prediction of chemo/immunotherapy response as well as the identification of mechanisms of resistance in metastatic lung cancer patients is still an unmet need[50].

In recent years microRNAs have emerged as master regulators of critical processes for lung cancer onset and progression[51]. Their role in driving lung cancer was found to be overall exerted through the expression regulation of targeted cancer-driver genes[51] and the modulation of complex cancer epigenetic mechanisms which impact tumor cells fitness by, for example, inducing EMT[52], stemness[53], immune evasion[54], and resistance to chemotherapy[55]. Furthermore, the exceptional stability of miRNA in harsh conditions and their presence in the body fluids[56] make them ideal candidates for the development of diagnostic, prognostic and predictive biomarkers [57,58].

Here, we performed a transcriptome analysis (miRNA and mRNA profiling) of LNmets of a cohort of patients with stage IIIA lung tumors by molecular profiling of EBUS and mediastinoscopy samples. We showed that N2 metastases resistant to NACT were characterized by an overall loss of miRNAs expression consistently with their prevalent role as tumor suppressors[59], as well as a profound reshape of the coding transcriptome. Our identified miRNA-based signatures (aka LN-signature) were accurate enough to predict NACT response which, to our knowledge, are the first of this kind and will warrant further investigations in larger and multicentric cohorts of patients.

Importantly, we unveiled that the miR-455-5p/PD-L1 axis regulates chemotherapy response of NSCLC cells, hallmarks metastases with active IFN-*γ* response pathway (an inducer of PD-L1 expression;[34]), and impacts T cells viability and relative abundances in TME (Fig. 7H). Remarkably, when we investigated the expression profile of miR-455-5p and correlated it with cisplatin sensitivity metrics, we found that loss of expression of miR-455-5p hallmarked intrinsic chemoresistance of NSCLC cell lines. This was in line with the miR-455-5p regulation in EBUS- and MED-samples which strongly suggested the relevance of miR-455-5p in controlling mechanisms of intrinsic and acquired chemoresistance. Indeed, we showed that miR-455-5p OE was sufficient to restore cisplatin sensitivity both *in vitro* and *in vivo*.

Several mechanisms involving drug accumulation, drug efflux and mediators of response to DNA damage have been implicated in platinum resistance so far[60]. Recently, PD-L1 was shown to regulate intracellular functions of cancer cells in a cell-autonomous way beside its immune-suppressive role on the membrane, including the regulation of cisplatin resistance [29,30]. NSCLC tumors treated with chemotherapy express higher levels of PD-L1 which, in turn, correlate with resistance and poor prognosis [26,27,61]. In keeping with this, we observed that PD-L1 expression is increased in resistant cells (both at basal level and upon cisplatin treatment) and direct inhibition of PD-L1 expression sensitize cells to cisplatin treatment. Importantly, we found that miR-455-5p directly targets PD-L1 in lung cancer cells and inhibits its expression thus contributing to response to cisplatin treatment. Intriguingly, other miRNAs of our LN-signature (i.e. miR-140-3p, miR-324-5p, miR-15b-5p and miR-93-5p) target PD-L1[62] which further enforces the role of PD-L1 in NACT response in stage IIIA patients.

miR-455-5p expression has been found dysregulated in several human malignancies including colon cancer, hepatic cancer, NSCLC, gastric cancer and prostate cancer [63–67]. Recently, a work by Chen et al. has reported that miR-455-5p is able to regulate cisplatin resistance in bladder cancer via the HOXA-AS3–miR-455-5p–Notch1 axis[68]. However, in our study, neither the *HOXA-AS3* nor the *NOTCH1* expressions were found modulated upon miR-455-5p OE *in vitro* or in N2 metastases (Fig. S11 A-B). As a matter of fact, we noticed that the miR-455-5p overexpression resulted in either minor or no effect on cisplatin sensitivity in low PD-L1 expressing cells, thus highlighting the role of PD-L1 as a central mediator of the miR-455-5p activity in the context of drug resistance in NSCLC. A recent study suggested that miR-455-5p could target PD-L1 3’UTR in hepatocellular carcinoma cells[69]. However, the validation of the miRNA binding site in the PD-L1 gene was carried out only in a unphysiological context (e.g. luciferase-based assay) and was not even confirmed in real-world cohort of patients. Moreover, no data were presented about the role of miR-455-5p/PD-L1 axis in the regulation of cisplatin response and cancer immune evasion.

The binding of tumor PD-L1 with the receptor PD-1 on T cells activates a signaling cascade that alters the T cell activity in many ways, including the inhibition of T cell proliferation and survival, cytokine production and other effector functions[36]. Therefore, we expect that miR-455-5p-PD-L1 axis may have also a role in a non-cell-autonomous way by regulating cancer immune-evasion in LNmets of stage IIIA patients. As a matter of fact, we showed that LNmets, which express low level of miR-455-5p, are characterized by a higher amount of both PD-L1 and PD-1 mRNA together with a trend of reduction of CD8 T cells, as we predicted *in silico* by CIBERSORTx analysis. Although an immunohistochemistry (IHC) analysis of LNmets to measure PD-L1, PD-1 and T cell markers was not feasible due to limited amount of samples, we showed in primary NSCLC tumors that higher level of miR-455-5p was associated with decreased PD-L1 expression and increase in CD8+ T cell infiltration, in line with our hypotheses. Recently, FDA approved neoadjuvant nivolumab plus p-doublet chemotherapy in resectable NSCLC regardless of PD-L1 tumor status[9]. Despite PD-L1 expression modulation was associated to immunotherapy response[70], PD-L1 has not been considered as reliable biomarkers mainly due to its spatial and temporal heterogenous expression[71] with PD-L1 negative tumors which responded also to ICIs[72]. However, GSEA analysis revealed that N2 metastases were enriched in a set of genes belonging to IFN-γ signature. IFN-γ is a proinflammatory cytokine produced by T cell and NK cells and is able to increase PD-L1 levels in cancer cells, thus promoting the inhibition of the T cell activity in the TME[73]. Moreover, IFN-γ-related gene signatures have been recently reported to predict the response to anti-PD-1 therapy in melanoma[74] and NSCLC patients[75]. Interestingly, our data indicate that miR-455-5p overexpression *in vitro* is able to decrease both IFN-γ-mediated PD-L1 expression and the enrichment in IFN-γ related genes observed in resistant cells, which deserves further investigations to explore the role of miR-455-5p and the overall LN-signature as potential reliable biomarkers to predict the response to ICIs. Moreover, given the ability of miRNA-based LN-signature to accurately predict NACT response, such signature could also be exploited in future studies as a potential biomarker for the newly approved drug regimen based on ICIs plus NACT.

Further studies have also highlighted a high tumor heterogeneity between metastatic lesions and primary tumors in the same NSCLC patients both in terms of pathway activation and PD-L1 expression[76], which may impact chemotherapy and immunotherapy response.

Although a direct comparison between nodal metastases and primary tumors was unfeasible in our cohorts, our data represent an important step forward in understanding the molecular mechanisms driving chemoresistance in lung cancer metastatic cells. Furthermore, we provided evidences for an unedited contribution of the miR-455-5p-PD-L1 axis in the regulation of chemoresistance and immunoevasion at the level of lymph nodal metastases, thus adding new grounds for bringing chemo-immunotherapy a step closer to stage IIIA clinical practice.

## CONCLUSIONS

Here, we showed that treatment naïve LNmets were characterized by distinct miRNA expression patterns which were predictive of NACT response. Importantly, by coupling whole-miRNA and mRNA profiling, we unveiled a key role for the miR-455-5p/PD-L1 axis which regulates chemotherapy response and immune evasion in metastatic NSCLC cells. To our knowledge, our study represents the most comprehensive transcriptome (coding and non-coding) analysis of LNmets in NSCLC patients. In conclusion, we described novel miRNA-based biomarkers and unveiled relevant mechanisms for LNmets resistance to chemotherapy which will contribute to improve outcome of lung cancer patients.

## METHODS

### Tumor sample collection and processing

#### EBUS samples

samples were obtained and processed as previously described[11]. EBUS-TBNA samples were collected from the mediastinal LNs station 4 and 7 of patients using a convex-probe (EBUS Convex Probe BF-UC180F; Olympus), a dedicated ultrasound processor (EU-ME2; Olympus) and a 22-gauge dedicated needle (Vizishot NA-201SX-4022; Olympus). One dedicated needle passage was put into cell culture medium for primary cell cultures expansion. Briefly, EBUS-TBNA samples were centrifuged for 5 min at 1,000 g at RT, resuspended in complete medium[11] and cultured on collagen-I rat tail (Gibco) coated plates for 6 to 12 days prior to total RNA extraction (Table 1). For long term expansion, primary cell cultures were expanded in Pneumacult Basal Ex (Stemcells technologies). EBUS cell line used for transfection experiments was derived from LNmets of a 54 years old female with lung adenocarcinoma. Criteria for selection of patients were: i) pathologically confirmed stage IIIA-pN2 NSCLC; ii) not having been treated before for their disease; iii) suitability for NACT followed by surgery.

#### MED samples

Two FFPE tissue sections (5–10µm thick) on glass slides with adequate tumor cellularity (>60%) were selected by a certified pathologist and microdissected by scraping with a scalpel prior to RNA isolation as previously described[11] (Table S2). Criteria for selection of patients were: i) pathologically confirmed stage IIIA-pN2 NSCLC; ii) not having been treated before for their disease; iii) suitability for NACT followed by surgery.

#### CIMA-CUN and CSS cohorts

tumor samples were obtained from NSCLC patients who underwent surgical resection at Clínica Universidad de Navarra (Pamplona, Spain) (CUN) and at the Casa Sollievo della Sofferenza Research Hospital (San Giovanni Rotondo, Italy) (CSS), respectively. Inclusion criteria were: i) absence of cancer within the previous five years; ii) complete resection of the primary tumor; iii) no adjuvant therapy prior to surgery. Tumors were classified according to the WHO 2004 classification and the 8^th^ TNM edition was used for tumor staging. RNA was extracted from one to two FFPE tissue sections (5µm thick) on glass slides with adequate tumor cellularity (>60%), selected by a pathologist. See also Table S5 and Table S6.

### Cell lines

NCI-H2023, NCI-H1993, NCI-H1975, NCI-H838, NCI-H1944, NCI-H1437, NCI-H1573, NCI-H2126, NCI-H322M, BEAS-2B and Jurkat were obtained from ATCC and cultured in RPMI (Gibco) with 5% FBS, 1% penicillin/streptomycin except for Jurkat medium, which was supplemented with 10% FBS. Primary cell cultures from LNmets of stage IIIA NSCLC were obtained and maintained as previously described[11]. All cell lines were grown at 37°C in a humidified incubator with 5% CO_2_ and routinely tested for Mycoplasma contamination using PCR.

### Creation of cisplatin resistant cells (CDDP-R)

Cisplatin (P4394, Sigma-Aldrich) was dissolved in vehicle solution (NaCl 0.9%) at a final concentration of 1 mg/ml and stored in the dark at RT for a maximum of 28 days. NCI-H2023 cells were subjected to treatment cycles (n=11), consisting of 3-4 days of cisplatin treatment and 1-2 weeks of culture in RPMI 5% FBS 1% penicillin/streptomycin to allow survived cells (i.e., the CDDP-R) to proliferate. The dose at the first treatment cycle was 0.6 µM then increased in subsequent cycles until reaching a maximum dose of 10 µM. Parental cells treated with vehicle solution were cultured in parallel and used as a control.

### Cell viability assay

Cells were seeded in 96-well plates in triplicate in 90 ul of complete media. At day 1 post seeding, cells were treated with increasing doses of cisplatin (3-fold serial dilution), or vehicle solution as a control. Cell viability was assessed by adding CyQUANT Cell Proliferation Assay Kit (Life Technologies) in a ratio of 1:10 directly in complete media. Fluorescence was measured at 480/528nm using a Sinergy HT (Biotek) microplate reader and IC_50_ was estimated using the online tool GR calculator[77].

### Cell transfection experiments

All transfection experiments were carried out by performing reverse transfection with Lipofectamine RNAiMAX (Thermofisher Scientific) according to the manufacturer’s instructions. The following oligos at the indicated concentration were used: 5nM of miR-455-5p mimic (MSY0003150, Qiagen) or recommended All Stars negative control siRNA (cat. 1027281, Qiagen); 7.5nM of PD-L1-specific miR-455-5p TSB (339194; sequence: GTAGACTATGTGCCTTTGCTCAG; Qiagen) or scramble TSB (339194; sequence: ACGTCTATACGCCCA; Qiagen); 10nM of siRNA against *CD274* (HSS120932, Thermofisher scientific) or recommended Stealth RNAi negative control Med GC (12935-300, Thermofisher scientific).

### Jurkat T cell apoptosis assay

Transfected NCI-H1975 were seeded overnight to allow them to adhere to culture plates. The day after, tumor cells were stimulated with 40ng/ml of IFN-γ for 8 hours and then co-cultured with Jurkat cells in the presence of Immunocult human CD3/CD28/CD2 T cell activator (Stemcells technologies) at a Jurkat cells to NCI-H1975 ratio of 1:4. After 72 hours, Jurkat cells were recovered from the co-culture and analyzed by AnnexinV-488 (Thermofisher) and 7-AAD (BD Pharmingen) staining in a BD FACS CANTO Cytometer. Gating strategy used to analyze apoptosis was reported in Fig. S7.

### Total RNA (including small RNA) isolation

Total RNA from commercial cell lines, EBUS samples and MED samples was isolated using respectively miRNeasy kit, AllPrep DNA/RNA/miRNA Universal Kit and AllPrep DNA/RNA FFPE Kit, respectively, according to the with manufacturer’s instructions. Total RNA quantification was carried out using the NanoDrop® ND-1000 spectrophotometer or Qubit RNA HS Assay Kit (Invitrogen).

### Quantitative Real Time-PCR (qRT-PCR) of miRNAs and mRNAs

For qRT-PCR of miRNAs, 10 ng of total RNA were reverse transcribed using a TaqMan MicroRNA Reverse Transcription Kit (ThermoFisher Scientific) and RT specific primers for miRNAs (ThermoFisher Scientific, See Table S9). 2.5 uL of RT product were pre-amplified for 14 cycles using the TaqMan PreAMP Mastemix and miRNA Taqman assay (see Table S9). The expression levels of miRNAs were normalized to the housekeeping gene U6 snRNA. For qRT-PCR of transcripts of commercially available cell lines, 1 ug of total RNA was reverse-transcribed using a High Capacity cDNA Reverse Transcription Kit (ThermoFisher Scientific) according to manufacturer instructions. For qRT-PCR of transcripts of EBUS and MED samples, 200 ng of total RNA were reverse transcribed with the SuperScript VILO cDNA Synthesis Kit (Thermo Fisher Scientific) in 20 µL of final volume and then cDNA was pre-amplified for 10 cycles. cDNA was amplified with the TaqMan Gene Expression assay (ThermoFisher Scientific, See Table S9) and QuantStudio 12k Flex thermocycler (ThermoFisher Scientific) using the manufacturer’s recommended cycling conditions. Data were normalized using the geometric mean of 3 genes (ESD1, GUSB and HPRT) as reference. Data normalization for both miRNAs and mRNAs was performed by using the delta-delta CT method or the calculation of the normalized Cq as previously described[78].

### Whole miRNA expression profile

10 ng of total RNA was reverse transcribed with MegaplexTM miRNA-specific stem-loop RT Primers Human Pool A v 2.1 (Thermo Fisher Scientific) and TaqMan® MicroRNA reverse transcription kit (Thermo Fisher Scientific) according to the manufacturer’s instructions. 5 µL of reverse transcribed product were pre-amplified for 14 cycles using the TaqMan PreAMP Mastemix and Megaplex PreAMP primers Pool A v 2.1 according to the manufacturer’s instructions (Thermo Fisher Scientific). The PCR reaction was performed using the TaqMan Universal Master Mix II, No AmpErase UNG (Thermo Fisher Scientific) by loading 100 µL of the pre-amplified mixture (final dilution 1:200) in each of the eight lanes of the TaqMan® Low Density Array miRNA Panel A v 2.0 (Thermo Fisher Scientific). Real-Time PCR was carried out on the QuantStudio 12k (Thermo Fisher Scientific) according to the manufacturer’s cycling conditions and by setting an automatic threshold. Cq data of miRNAs were normalized (Cqn) using U6 snRNA as previously described[78]. miRNAs with a Cq<30.01 in at least 50% of samples among one of the experimental groups tested in the analysis, were considered as detected.

### Zebrafish cell-derived xenograft (zCDX)

zCDX models were developed by a CRO (BioReperia AB). Transgenic Tg(fli1:EGFP)y1 zebrafish embryos were raised at 28°C for 48 hours in E3 embryo medium (containing per liter: 0.286g NaCl, 0.048g CaCl2, 0.081g MgSO4 and 0.0126g KCl with 0.2 mM 1-Phenyl-2-Thiourea aka PTU). At 2 days post fertilization, embryos were injected subcutaneously in the perivitelline space with transfected parental and CDDP-R cells previously labeled with FAST Dil^TM^ oil (ThermoFisher Scientific) and treated with ± Cisplatin 5 mg/L for 3 days. Images of tumors were taken by using a fluorescent stereoscope with a K5 camera (Leica) and LAS X software v3.7.1.21655 at 100x magnification with no binning. Images of tumors were taken right after injection (day 1) and after drug treatment (day 4). Images were automatically analyzed by using the HuginMunin software v2.7.0.0 (BioReperia AB). Tumor growth regression was calculated by dividing the number of tumor pixels at day 4 by the number of tumor pixels at day 1 in the same embryo and multiplied by 100.

### Genome-wide expression profiling

Gene expression profiling of MED samples and NSCLC cell lines (two independent biological replicates) was carried out using the GeneChip® Pico reagent Kit and the GeneChip® WT Plus reagent Kit, respectively. For both reagents, the GeneChip® Human Clarion S Array (Thermofisher Scientific) was used according to the manufacturer’s instructions. Quality control, normalization of CEL files and statistical analysis were performed using the Transcriptome Analysis Console (TAC) software v4.0 (Thermo Fisher Scientific) by performing the “Gene level SST-RMA” summarization method with human genome version hg38. Differentially expressed genes were defined as those with a fold-change (FC) difference of at least 1.5 and a p-value less than 0.05. For MED samples, 5 pN2 and 5 pN0 samples balanced for age, sex and histotype were pooled to obtain 2 pools for each experimental condition (pN2 and pN0). Microarray expression data can be found at GEO database (GSE193707; reviewer token: qtsheiycfpsvxsr).

### Predictive risk model

A ridge-penalized unconditional logistic regression was applied in the training set to model the odds of responding as function of the 16-miRNAs that were scored as differentially expressed between responder and non-responder patients in the EBUS-samples (16 miRNA model). The same strategy was used for the 14-miRNA and 4-miRNA models. Cross-validated (10-fold) log-likelihood with optimization (50 simulations) of the tuning penalty parameter was applied. Probability of being responder was estimated, and model performance was assessed using the area under the receiver operating curve (AUC). Min-max scaling of miRNAs expression in the validation set was implemented before applying the predictive model. LASSO approach was used to reduce the number of predictors.

### Analysis of cell line publicly available datasets

Cell viability of cisplatin for the indicated dataset was downloaded directly from the Depmap portal (https://depmap.org/portal/compound/cisplatin?tab=dose-curves). Analysis of cell viability data was restricted only to NSCLC cell lines for which miRNA expression data was available in the CCLE dataset. Median cell viability was calculated at each concentration and plotted. Quality control (QC) for IC_50_ estimation was applied following instructions reported in Sebaugh et al.[79]. Briefly, we estimate IC_50_ values for cell lines in each dataset by taking advantage of cell viability data downloaded from Depmap portal and the online software ‘GR calculator’. QC criteria applied were: at least two concentrations below the 50% response concentration and above the 50% response. Only proportions of cell lines in each dataset for which IC_50_ estimation was accurate according to Sebaugh et al. were reported (see Table S2).

### CIBERSORTx analysis

CIBERSORTx[44] was run using the online web-tool (https://cibersortx.stanford.edu) and following the developers’ instructions. The CIBERSORTx analysis was conducted using the following settings: LM 22 as signature matrix file, absolute mode running and 100 permutations. CIBERSORTx score is an estimation of cell fraction of each specific subpopulation in each tumor sample. CIBERSORTx complete results were reported in Table S7.

### Gene Set Enrichment Analysis (GSEA)

GSEA (GSEA, https://www.gsea-msigdb.org/gsea/index.jsp) was performed using Signal2Noise metric, 1000 random sample sets permutation, and median gene expression values for class comparison. For enrichment analysis of hallmarks of cancer, we used the gene matrix h.all.v7.4 symbols.gmt available from MSigDB. For miR-455-5p target enrichment analysis, we built a custom gene matrix by including human genes that were highly or moderately predicted to be miR-455-5p targets (cumulative weighted context++ score<=-0.2) by Target Scan (release 7.2) and were well expressed (log_2_ intensity>4) in all samples used in each analysis. Significant gene sets were considered as those with a false-discovery rate (q-value) less then 5%. For single-sample gene set enrichment analysis of TGCA cohorts, ssGSEA scores were calculated by using the GSVA package in R. Gene signature for exhausted CD8+ T cell were obtained from Cai et al.[42] while gene signatures for IFN-γ and IFN-α response were downloaded from MSigDB hallmark gene sets (version h.all.v7.4 symbols.gmt).

### Statistics

Hierarchical clustering was performed using Cluster 3.0 (C Clustering Library 1.56; http://bonsai.hgc.jp/~mdehoon/software/cluster/software.htm) and Java Tree View (Version 1.1.6r4; http://jtreeview.sourceforge.net). The uncentered correlation and centroid linkage clustering method was used. Heatmaps were obtained by using MORPHEUS (https://software.broadinstitute.org/morpheus/) or Java Tree View. All graphs and statistical analyses were performed using Prism (version 7.0e), SPSS (version 15.0), SAS software and R 3.3.1 (R Core Team, 2016). The normality of data was controlled by Shapiro-Wilk and D’Agostino & Pearson normality tests. The details about statistical tests, number of independent replicates (N) and definition of error bars were specified in the Fig. legends. Statistical output (p-value) was represented by asterisks as follows: non-significant (ns) > 0.05, \**p* ≤ 0.05, \*\**p* ≤ 0.01, \*\*\**p* ≤ 0.001, \*\*\*\**p* ≤ 0.0001. A p<0.05 was considered to be statistically significant. Sample size for tissue-based assays was chosen on the basis of sample availability. The investigators were not blinded when analyzing the data except for IHC analysis and zebrafish experiments.

## LIST OF ABBREVIATIONS

3’-UTR: 3’ untranslated region
ACTA2: Actin Alpha 2, Smooth Muscle
AUC: Area under the curve
CC10: Clara cell 10
CCLE: Cancer Cell line Encyclopedia
CD90: Cluster of Differentiation 90
CDDP: Cisplatin
CDH5: Cadherin 5
CTRL: Control
CTRPv2: Cancer Therapeutics Response Portal
DMSO: Dimethylsulfoxide
E-CAD: E-cadherin
EBUS-TBNA: endobronchial ultrasound transbronchial needle aspiration
EGFR: Epidermal Growth Factor Receptor
EMT: Epithelial-to-mesenchymal transition ESD1: Esterase D
FDR: False discovery rate
FFPE: Formalin fixed paraffin embedded
GAPDH: Glyceraldehyde-3-Phosphate Dehydrogenase
GDSC1-2: Genomics of Drug Sensitivity in Cancer
GET: Gene signature of exhausted CD8+ T cell
GSEA: Gene set enrichment analysis
HPRT1: Hypoxanthine Phosphoribosyltransferase 1
ICI: Immune checkpoint inhibitors
IFN-α: Interferon-alpha
IFN-*γ*: Interferon-gamma
IHC: Immunohistochemistry
KRT5: Keratin 5
LN: Lymph node
LNmets: Lymph nodal metastases
LUAD: Lung adenocarcinoma
LUSC: Lung squamous cell carcinoma
MED: Mediastinoscopy
N-CAD: N-cadherin
NACT: Neoadjuvant chemotherapy
NES: Normalized enriched score
NK: Natural Killer
NKX2-1: NK2 Homeobox 1
NSCLC: non–small-cell lung cancer
OE: Overexpression
P-doublet: Platinum-based doublet
Pan-CK: pan-cytokeratin
PRISM: Profiling Relative Inhibition Simultaneously in Mixtures
PTPRC: Protein Tyrosine Phosphatase Receptor Type C
SFTPC: Surfactant Protein C
SLUG: Zinc finger protein SNAI2
SOX2: SRY-Box Transcription Factor 2
TILs: Tumor-infiltrating lymphocytes
TLDA: TaqMan Low-density Array
TME: Tumor-microenviroment
TSB: Target site blockers
TWIST1: Twist Family BHLH Transcription Factor 1
VIM: Vimentin
zCDX: Zebrafish cell derived xenograft
ZEB1: Zinc Finger E-Box Binding Homeobox 1

## DECLARATIONS

### Ethics approval and consent to participate

The institutional ethical committees approved this study (registration number: R65/14-IEO76; BIO-EBUS-V1.0_Ott19; BIO-POLMONE-V1.0_Giu16), and informed consent was obtained from all patients enrolled.

### Consent for publication

Not applicable

### Availability of supporting data

All data generated or analysed during this study are included in this published article and its supplementary information files. The datasets generated during the current study (microarray expression data) are available at GEO database (GSE193707; reviewer token: qtsheiycfpsvxsr). The web link to public datasets analysed during the current study are available in the materials section.

### Author’s contribution

Conceptualization: RC, FB Methodology: RC, TC, ED, VM, OP, EB, FM, MB, PG, LM, CC, JG, LS, JS, AP, FB Investigation: RC, TC, ED, MC, FMZ, LM, JG, LS, FB Visualization: RC, TC, ED, VM, LM, FB Funding acquisition: FB Project administration: FB Supervision: RC, FB Writing – original draft: RC, FB Writing – review & editing: RC, FB

## Acknowledgement

We thank Dr. Lucia Anna Muscarella and Dr. Vincenzo Giambra (Fondazione IRCCS Casa Sollievo della Sofferenza; San Giovanni Rotondo, Italy) for sharing NCI-H1573, NCI-H2126, NCI-H1975 and Jurkat cells respectively. We thank Dr. Karmele Valencia for her help in performing IHC analysis of CIMA-CUN cohort and Dr. Rose Mary Carletti for technical support with TLDA qRT-PCR experiments.

We thank Dr. Miriam Kuku Afanga for technical support with immunoblot experiments. Dr. Fabrizio Bianchi wishes to thank Prof. Pier Paolo di Fiore (European Institute of Oncology) for his support and mentorship. We are grateful to Chiara Di Giorgio for english editing and manuscript proofreading.

