## Supplementary Materials for "miRNome profiling of lung cancer metastases revealed a key role for miRNA-PD-L1 axis in the modulation of chemotherapy response"

**SUPPLEMENTAL MATERIALS**

**Cell Proliferation Assay**

Cell proliferation was evaluated with the CyQUANT Cell Proliferation Assay Kit (Life Technologies), according to the manufacturer’s instructions. Briefly, cells were seeded into 96-well plates in 90μl of complete media at the previously determined seeding density and CyQuant was added directly in cell media at 24h and 96h. Fluorescence was measured using a microplate spectrofluorometer (Synergy HTX; BioTek) at 480/520 nm, respectively. The doubling time was calculated as follows:

$$doubling time=\frac{T*ln \left( 2 \right)}{log\left( \frac{X\left( 96h \right)}{X\left( 24 h \right)} \right)}$$

Where:

- *X(96h)* = number of cells at 96 hours
- *X(24h)* = number of cells at 24 hours
- *T* = time in hours (i.e., 72 hours)

**Cell migration assay**

Cells were seeded onto 35mm dish containing 4 well-culture insert (Ibidi) in complete medium. After appropriate cell attachment (24 hours), cells were starved for 12h in RPMI with 1% FBS. After the removal of culture-insert, cells were washed twice with 1X PBS and cultured in RPMI with 10% FBS. Cell migration was evaluated at the indicated time points with a 10X objective using an Eclipse TE300 fluorescence microscope (Nikon). The cell free area (percentage of control) at indicated time points was measured by Fiji software (ImageJ; https://imagej.nih.gov/ij/).

**Cell invasion assay**

Invasion assays were performed by using trans-well chambers with 8.0‐µm pore polycarbonate membrane (Costar, Corning Inc., Corning, NY, USA) inserted in a 24-well plate and coated with 200ul of Corning® Matrigel® matrix at a concentration of 150 µg/ml for 24 hours at 37°C. 1x10^5^ cells were seeded in the upper chamber in RPMI free medium while the lower chamber was filled with RPMI with 20%FBS as chemoattrantant. After 24h of incubation at 37°C in a humidified incubator with 5% CO_2_, non-invading cells remained in the upper chamber were mechanically removed by using cotton swabs and washed away with 1XPBS while those on lower surface (invading cells) were fixed using 4% PFA and cells nuclei stained with DAPI (Sigma-Aldrich). Images of ten fields for each experiment were acquired at 10X magnification using Eclipse TE300 Microscope (Nikon). Invaded cells were counted (nuclei) with Cell Count tool of Fiji software (ImageJ) and the invasion rate was calculated as the number of invading cells divided by the total number of cells seeded.

**Western blotting**

Cells were lysed for ten minutes in a modified Laemmli sample buffer (2% SDS, 20% glycerol, and 125 mM Tris– HCl, pH 6.8) at 100°C, then quantified using the BCA Protein Assay Kit (Pierce). Equal amounts of proteins were separated by SDS–PAGE (Mini-PROTEAN® TGX TM, BioRad) then transferred on a polyvinylidene fluoride (PVDF) membrane (Bio-Rad) using Trans-blot Turbo (Bio-Rad). PVDF membranes were incubated with primary (O/N at 4°C) (see Table S8) and HRP-linked secondary antibodies (1h at RT) (see Table S8) after 1h of blocking at RT with either 5% BSA or 5% non-fat dry milk in Tris-buffered saline with 0.1% Tween-20 (TBS-T). After incubation with chemiluminescent substrate (ClarityTM Western ECL Substrate, BioRad), specific signals were acquired using ChemiDoc XRS gel imaging system (Bio-Rad). Densitometric analysis of WB bands was performed with ImageLab Software (version 5.2.1) (Biorad) and normalized on GAPDH (loading control). Data quantification was reported in the respective figures as well as in Data File 4. One sample t test was used to calculate statistical significance.

**Immunofluorescence**

Cells were seeded in 15μ-slides (Ibidi) and after 3 days fixed for 10 minutes at RT in a 4% solution of Paraformaldehyde (PAF). The cells were permeabilized for 5 minutes with 0.05% TritonX-100 in PBS and subsequently incubated for 1 hour at RT with a blocking solution (PBS with 5% donkey serum and 2% BSA). After blocking, cells were incubated with primary antibody for 1 hour at RT (see Table S8) and then with a secondary antibody conjugated to a fluorophore (see Table S8) for 2 hours at RT. Cells were then counterstained with DAPI and mounted with Vectashield anti-fade solution (Vector Laboratories). The staining analysis was carried out using a LEICA SP8 confocal microscope using the 63x oil immersion objective. The images were acquired using Leica Confocal Software and then edited with Adobe Photoshop. The adjustments used in the preparation of the figures were for brightness, contrast, and background noise (blur filter). For comparison purposes, images of the same antigen were acquired under constant acquisition settings and equal post-acquisition adjustments.

**Flow cytometry**

1x10^5^ cells were collected, resuspended in FACS staining solution (PBS 1x with 3% FBS) and then stained for 1h at 4°C in the dark with primary antibodies conjugated to a fluorophore (see Table S8). DAPI (Sigma) was used to identify dead cells during the analysis. FACS analysis was performed using a Beckman Coulter MoFlo Astrios or a BD FACS Canto and data processed with FlowJo software (v.10). MFI indicates median fluorescence intensity except for Figure S5 C to E where we used MFI as mean fluorescence intensity to perform correlation with qRT-PCR and Western blot Data.

**IHC analysis**

IHC analysis was performed by an automated Autostainer Link 48 (Agilent Technologies) platform according to the manufacturer’s instructions. The following antibodies were used: anti-human PD-L1 monoclonal antibody (22C3 clone PharmDx; Agilent DAKO) and anti-human anti-CD8 (clone C8/144B; Agilent).

**Analysis of gene/miRNA expression profile from public available dataset**

miRNA and gene expression data of TGCA-LUAD and TGCA-LUSC cohort were downloaded from the cBIO data portal (<https://www.cbioportal.org/>). Clinical data from both cohorts were retrieved from GDC portal (<https://portal.gdc.cancer.gov/>). For the analysis of the association of miR-455-5p expression and clinical-pathological features, all samples with miRNA data available were included in the analysis (N_LUAD_=510 and N_LUSC_=478). For correlation analysis, we extracted only those samples for which both miRNA and gene expression data were available, resulting in a total of 507 LUAD tumors and 473 LUSC tumors. The dataset was then stratified based on miR-455-5p expression in tertiles (high [top tertile], int [middle tertile] and low [low tertile] to perform correlation analysis subcategory with the same number of patients and with a different degree of expression of miR-455-5p.

For GSE33072 dataset, gene expression data (log2-RMA) of core biopsies from chemorefractory NSCLC patients (N=131) were downloaded from the GEO Database (<https://www.ncbi.nlm.nih.gov/geo/>).

**SUPPLEMENTAL FIGURES LEGENDS**

**Fig. S1. Whole miRNA expression profile of chemo-naïve metastatic tissue from stage III NSCLC patients.** (**A**) Representative H&E images of biopsies of mediastinal lymph nodes before (upper) and after (lower) microdissection of tumor cells. Scale bar: 300 μm. (**B**) Number and percentage of miRNAs detected (yellow) or not detected (blue) in MED-cohort. (**C**) Number and percentage of miRNAs detected in both MED- and EBUS-cohort (Common), only MED-cohort or only EBUS-cohort. (**D**) Violin plot representing the expression (Cqn) of all miRNAs detected in MED-cohort in EBUS-cohort. (**E**) Dot plot representing miRNA fold changes in chemoresistant (pN2) vs. chemosensitive (pN0) samples in MED- and EBUS-cohort. Each dot represents a single miRNA. Only detected miRNAs were plotted. Red bar identifies the median. P-value testing the difference in the proportion of downregulated miRNAs versus 50% was calculated by Binomial proportion test. (**F**) Volcano plot representing differentially expressed miRNAs in chemoresistant (pN2) vs. chemosensitive (pN0) metastatic lung tumor cells collected by MED before NACT as reported in (A). P-value was calculated using the Mann-Whitney U test. Grey dot, unchanged; Blue dot, downregulated (FC>|1.5|; p-value<0.05); Red dot, upregulated (FC>|1.5|; p-value<0.05). (**G**) Venn diagram representing the overlap of differentially expressed miRNAs (FC>|1.5|; p-value<0.05) in EBUS or in MED cohorts. P-value was calculated using Exact hypergeometric probability with normal approximation.

**Fig. S2. Biological characterization of Cisplatin resistant cells.** (**A**) Dose-response curves of Parental and CDDP-R cells treated with cisplatin for 72 hours. Error bars indicate SEM (N=4). (**B**) Bar plot representing cisplatin potency (IC_50_, left) and efficacy (E_max_, right) of Parental and CDDP-R cells. The result is shown as fold change relative to Parental cells. Data represent mean ± SEM (N=4). P-values were calculated by one sample t-test. (**C**) Representative brightfield images of Parental and CDDP-R cells. Scale bar: 100µM. (**D**) WB analysis of EMT markers in Parental and CDDP-R cells. GAPDH was used as loading control. These data are representative of three independent experiments. (**E**) Representative images of confocal analysis of E-cadherin (E-CAD; red) and Vimentin (VIM; red) expression in Parental and CDDP-R cells. DAPI (light blue) visualizes nuclei. Scale bar: 50μm. (**F**) qRT-PCR analysis of expression of EMT markers in Parental and CDDP-R cells. Data are mean ±SEM (N=3). Fold change is relative to Parental cells condition. P-value was calculated by one sample t-test. *P<0.05. (**G**) Volcano plot showing differentially expressed genes found by microarray analysis between CDDP-R CTRL (N=2) vs Parental CTRL (N=2). Grey dot, unchanged genes; Blue dot, downregulated genes (p-value <0.05; FC >|1.5|); Red dot, upregulated genes (p-value <0.05; FC >|1.5|). The number of significantly regulated genes is shown. P-value was calculated using the Limma moderated t-test. (**H**) GSEA of EMT gene sets in CDDP-R CTRL vs Parental CTRL cells. NES normalized enrichment score, FDR false-discovery rate based on 1000 random samples permutations. (**I**) Lollipop plot showing GSEA results using the ‘Hallmark gene sets’ collection in CDDP-R CTRL vs Parental CTRL cells. (**J**) Doubling time (hours) of Parental and CDDP-R cells. Data are mean ± SEM (N=4). P-value was calculated by performing t-test with Welch’s correction. (**K**) Wound healing assay of Parental and CDDP-R cells. Left: representative brightfield images of migrating Parental and CDDP-R cells at the indicated time points. The yellow line highlights the borders of the wound. Scale bar=100µM. Right: bar plot showing the percentage of cell free area compared to the initial scratch area (N=4). Statistical significance was computed using t-test with Welch’s correction. (**L**) Cell invasion assay of Parental and CDDP-R cells. Left: representative fluorescence image of invading Parental and CDDP-R cells at 24 hours post seeding. DAPI (light blue) visualizes nuclei. Scale Bar: 100µM. Right: bar plot representing the invasion rate of Parental and CDDP-R cells. The result is shown as fold change relative to Parental cells. Data represent mean ± SEM (n=3). P-value was calculated by one sample t-test.

**Fig. S3. Implantation analysis of CDX in zebrafish.** (**A**) qRT-PCR of miR-455-5p expression in Parental CTRL, CDDP-R CTRL and NCI-H2023 CDDP-R OE cell lines before injection in the perivitelline space of zebrafish larvae. Data, expressed as Cqn, are mean ± SD (N=3). (**B**) Representative fluorescence images of tumor masses at day 1 (implantation). Dil (red) identifies tumor cells. Scale bar: 200μm. (**C**) Quantification of the size of tumor masses at day of implantation. Columns represents mean ± SEM (N=36-38, for each cell line). Results are expressed as tumor area (pixel density). Each dot identifies individual zebrafish larvae. P-value was calculated using the Mann-Whitney U test. *P<0.05, **P<0.01; ns, not significant.

**Fig. S4. miR-455-5p overexpression decreases the proliferation rate of NSCLC cells. (A** and **B)** Doubling time (hours) of NCI-H1993 (A), Parental (B) and CDDP-R cells (B) transfected with miR-455-5p (OE) or mimic negative control (CTRL). Data are mean ± SEM (N=4). P-value was calculated by performing a t-test with Welch’s correction.

**Fig. S5. Higher basal levels of PD-L1 are associated to cisplatin resistance in NSCLC *in vitro*.** (**A**) qRT-PCR analysis of *CD274* in the indicated cell lines. Data, expressed as Cqn, are mean ± SD of three technical replicates. (**B**) Immunoblot analysis of PD-L1 expression in the indicated cell lines. GAPDH was used as loading control. Fold change (FC) is relative to NCI-H1993 cells. (**C**) Flow cytometry analysis of PD-L1 expression on cell-surface in the indicated cell lines. Histograms represent PD-L1 fluorescent intensity (x-axis) versus the number of events relative to mode (% of Max, y-axis). Mean fluorescence intensity is also reported. (**D**) Distribution of cisplatin potency (IC_50_) and efficacy (E_max_) in relation to PD-L1 mRNA (left panel), total protein (central panel) and cell-surface protein (right panel). Color of the dots indicates the amount of PD-L1. (**E**) Correlation matrix between PD-L1 mRNA, total (tot) and cell-surface (c-surf) PD-L1 protein and cisplatin sensitivity metrics (IC_50_ and E_max_). Color and size of the bubbles indicate the correlation coefficient R and the P-value, respectively. Statistical significance was computed by the Spearman correlation test. (**F**) Representative flow cytometry histogram plot (left) and quantification (right) of cell surface PD-L1 median fluorescence intensity (MFI) in NCI-H1993 cells transfected with a siRNA against PD-L1 or a scramble oligo as control. Data are presented as mean ± SEM (N=3). P-value was computed by one sample t-test. (**G**) Dose-response curves of NCI-H1993 cells transfected as in (F) and treated with cisplatin for 72 hours. Error bars indicate SEM (N=4). (**H**) Bar plot representing the cisplatin potency and efficacy of NCI-H1993si CTRL and NCI-H1993 siPD-L1. The result is shown as fold change relative to NCI-H1993 siCTRL cells. Data are mean ± SEM (N=4). P-value was computed by one sample t-test. ***P<0.001, **P<0.01

**Fig. S6. PD-L1 expression contributes to cisplatin resistance in an *in vitro* model of acquired resistance.** (**A**) qRT-PCR analysis of PD-L1 expression (mRNA) in Parental and CDDP-R cells. Data are expressed as fold change relative to Parental cells. Columns represent mean ± SEM (N=3). P-value was calculated by one sample t-test. (**B**) Immunoblot analysis of PD-L1 expression in Parental and CDDP-R cells. GAPDH was used as loading control. Experiments were performed in duplicates. (**C**) Flow cytometry of PD-L1 expression on cell-surface in Parental and CDDP-R cells. On the left panel, representative flow cytometry histogram plot of cell surface PD-L1 median fluorescence intensity (MFI) at the indicated experimental conditions (N=4 replicates). On the right, bar plot of PD-L1 expression on cell-surface (left panel) and percentage of PD-L1+ cells (right panel) in the indicated cell lines. The result is shown as fold change relative to Parental cells. Data are mean ± SEM (N=4). P-value were calculated by one sample t-test. (**D**) Representative flow cytometry histogram plots (left) and quantification (right) of cell surface PD-L1 MFI in Parental and CDDP-R cells treated with a siRNA against PD-L1 or a scramble oligo. Data are expressed as fold change in MFI relative to Parental cells treated with a scramble siRNA. Data are mean ± SEM (N=3). P-values were calculated by one sample-t test. *P<0.05, ***P<0.0001. (**E**) Dose-response of curve of Parental and CDDP-R cells transfected as in (D) and treated with cisplatin for 72 hours. Error bars indicate ± SEM (N=4). (**F**) Bar plot representing cisplatin potency and efficacy in Parental siCTRL, Parental siPD-L1, CDDP-R siCTRL and CDDP-R siPD-L1 cells. The results are shown as fold change relative to Parental siCTRL. Data are mean ± SEM (N=4). P-values were calculated by one sample-t test. * P<0.05, **P<0.01; ns, not significant.

**Fig. S7. Gating strategy used to analyze Jurkat T cells apoptosis after co-culture with NCI-H1975 tumor cells.**

Representative dot plots of doublet exclusion (left panel), gate of Jurkat T (central panel) and analysis (right panel) of apoptosis (Annexin V) and cell viability markers (7AAD) used in experiments presented in Figure 7C.

**Fig. S8. Analysis of miR-455-5p and PD-L1 association in post-chemotherapy samples of NSCLC patients.** (**A**) GSEA using miR-455-5p predicted target genes, GET signature and IRF4 signature in PD-L1 (aka *CD274*) high vs. PD-L1 low chemoresistant samples from the GSE33072 dataset. NES, normalized enrichment score; FDR, false-discovery rate. (**B**) Spearman correlation analysis of PD-L1 (aka *CD274*) and COL27A1 in chemoresistant samples from the GSE33072 dataset (N=131) **(C)** Heat map showing gene expression of PD-L1 (aka *CD274*) and COL27A1 in chemoresistant samples from the GSE33072 dataset (N=131).

**Fig. S9 miR-455-5p regulates IRF2 expression in NSCLC cell lines.** (**A-B**) Immunoblot analysis of IRF2 and PD-L1 in CDDP-R cells (A) and NCI-H1975 (B) transfected with a miR-455-5p mimic or a negative control. GAPDH was used as loading control.  **(C)** Schematic model of miR-455-5p-dependent regulation of PD-L1 and IRF2 in NSCLC.

**Fig. S10. Estimation of immune features of chemo-naïve lung metastatic tumor tissue. (A**) Estimation of “NK cell activated” and “T cell CD8” immune subpopulations in pN2 and pN0 metastasis samples obtained by CIBERSORTX deconvolution of transcriptomic profiles. Data are mean ± SEM (N=2). Each pool is composed by 5 individual samples balanced for age, gender and histotype. Statistical significance was indicated in the graph and was calculated by unpaired t test (**B**) Expression levels of MHC and immunomodulatory proteins for pN2 and pN0 metastasis. Data are log_2_-ratio. The size of the bubble is proportional to statistical significance while colors indicate fold change as per the legend.

**Fig. S11. miR-455-5p is not involved in the regulation of *NOTCH-1* and *HOXA-AS3* expression in NSCLC.** (**A to C**) Expression levels of *NOTCH1* and *HOXA-AS3* (Log_2_ intensity; microarray) in MED pN2 and pN0 (A), NCI-H1993 CTRL and OE cells (B) and CDDP-R CTRL and OE cells (C). Error bars represent SEM. Statistical significance was calculated by Limma moderated t-test. ns, not significant.

**SUPPLEMENTAL TABLES LEGENDS**

**Table S1. miRNA coefficients, area under the curve (AUC), receiver operating characteristic (ROC) curves, and box plot of the predicted probability of being a responder for the predictive models (Training).**

**Table S2. Quality control of IC50 estimation from publicly available dataset.** Abbreviation: NSCLC, non-small cell lung cancer; QC, Quality control; CV, coefficient of variation

**Table S3. GSEA of hallmark of cancer signatures of transcriptomic data.**

**Table S4. Clinical-pathological features of TGCA-LUAD and TGCA-LUSC cohorts.** Clinical information of patients from TGCA-LUAD and LUSC cohort stratified in tertiles based on miR-455-5p expression (see Methods). “High” and “low” tertiles were reported in the table. the Percentages could not add up 100 due to rounding; (a) median age of the combined LUAD and LUSC cohorts=67 years; (b) Fisher’s exact test. Significant P-value were highlighted in bold.

**Table S5. Clinical-pathological features of PD-L1 CSS and CIMA-CUN cohorts.** Clinical information and experimental data of patients from PD-L1 CSS cohort (N=8) (**A**) and CIMA-CUN cohort (N=7) (**B**). Experimental data were rounded to the second decimal place. Abbreviations: LUAD, Lung adenocarcinoma; LUSC, Lung squamous cell carcinoma; TPS, tumor proportion score; Cqn, normalized Cq.

**Table S6. Clinical-pathological features of CD8 CIMA-CUN cohort (A) and CD8-CSS cohort (B).** Clinical information and experimental data of patients from CD8-CIMA-CUN cohort (N=22) and CD8-CSS cohort (N=25). Patients were stratified in “high” and “low” based on the median value of the percentage of CD8+ cells. Percentage of CD8+ cells and miR-455-5p levels are expressed as z-score. Abbreviations: LUAD, Lung adenocarcinoma; LUSC, Lung squamous cell carcinoma; Cqn, normalized Cq.

**Table S7. Immune cell estimation of MED samples by CIBERSORTx.** Estimation of 22 immune cell populations obtained by deconvoluting transcriptomic data of chemoresistant (pN2) and chemosensitive (pN0) MED samples (N=2 pools for each experimental condition) using CIBERSORTx algorithm. Absolute values for each cell population was reported. Each pool is composed by 5 individual samples balanced for age, gender and subtype.

**Table S8. Primary and secondary antibody used for western blot, immunofluorescence and flow cytometry.**

**Table S9. Primer used for qRT-PCR of genes and miRNA.**
